# Bi-fated tendon-to-bone attachment cells are regulated by shared enhancers and KLF transcription factors

**DOI:** 10.1101/2020.01.29.924654

**Authors:** Shiri Kult, Tsviya Olender, Marco Osterwalder, Sharon Krief, Ronnie Blecher-Gonen, Shani Ben-Moshe, Lydia Farack, Hadas Keren-Shaul, Dena Leshkowitz, Tomer Meir Salame, Terence D. Capellini, Shalev Itzkovitz, Ido Amit, Axel Visel, Elazar Zelzer

**Affiliations:** Department of Molecular Genetics, Weizmann Institute of Science, Rehovot 76100, Israel; Environmental Genomics and Systems Biology Division, Lawrence Berkeley National Laboratory, Berkeley, CA, 94720, USA; Department of Immunology, Weizmann Institute of Science, Rehovot 76100, Israel; Department of Molecular Cell Biology, Weizmann Institute of Science, Rehovot 76100, Israel; Life Sciences Core Facilities, Weizmann Institute of Science, Rehovot 76100, Israel; Department of Human Evolutionary Biology, Harvard University, Cambridge, MA, USA; Broad Institute of Harvard and MIT, Cambridge, MA, USA; U.S. Department of Energy Joint Genome Institute, Lawrence Berkeley National Laboratory, Berkeley, CA, 94720, USA; School of Natural Sciences, University of California, Merced, CA 95343, USA

**Keywords:** Musculoskeletal system, Cartilage, Tendon, Enthesis, *Klf2*, *Klf4*, RNA-Seq, ATAC-Seq, Transcriptome, Mouse

## Abstract

The connection between different tissues is vital for the development and function of any organs and systems. In the musculoskeletal system, the attachment of elastic tendons to stiff bones poses a mechanical challenge that is solved by the formation of a transitional tissue, which allows the transfer of muscle forces to the skeleton without tearing. Here, we show that tendon-to-bone attachment cells are bi-fated, activating a mixture of chondrocyte and tenocyte transcriptomes, which is regulated by sharing regulatory elements with these cells and by Krüppel-like factors transcription factors (KLF).

To uncover the molecular identity of attachment cells, we first applied high-throughput RNA sequencing to murine humeral attachment cells. The results, which were validated by *in situ* hybridization and single-molecule *in situ* hybridization, reveal that attachment cells express hundreds of chondrogenic and tenogenic genes. In search for the underlying mechanism allowing these cells to express these genes, we performed ATAC sequencing and found that attachment cells share a significant fraction of accessible intergenic chromatin areas with either tenocytes or chondrocytes. Epigenomic analysis further revealed transcriptional enhancer signatures for the majority of these regions. We then examined a subset of these regions using transgenic mouse enhancer reporter. Results verified the shared activity of some of these enhancers, supporting the possibility that the transcriptome of attachment cells is regulated by enhancers with shared activities in tenocytes or chondrocytes. Finally, integrative chromatin and motif analyses, as well as the transcriptome data, indicated that KLFs are regulators of attachment cells. Indeed, blocking the expression of *Klf2* and *Klf4* in the developing limb mesenchyme led to abnormal differentiation of attachment cells, establishing these factors as key regulators of the fate of these cells.

In summary, our findings show how the molecular identity of bi-fated attachment cells enables the formation of the unique transitional tissue that connect tendon to bone. More broadly, we show how mixing the transcriptomes of two cell types through shared enhancers and a dedicated set of transcription factors can lead to the formation of a new cell fate that connects them.

## Introduction

The function of the musculoskeletal system relies on the proper assemblage of its components, namely skeletal tissues (bone, cartilage, and joints), muscles and tendons. However, the attachment of tissues composed of materials with large differences in their mechanical properties is highly challenging. In the musculoskeleton, elastic tendons, which have a Young’s modulus (a measure of stiffness) in the order of 200 megapascal, are attached to the much harder bone, with a modulus in the order of 20 gigapascal. This disparity makes the connection between these two tissues a mechanical weak point, which is subject to higher incidence of tearing by both external and internal forces acting on the musculoskeleton during movement. The evolutionary solution to this problem is the enthesis, a transitional tissue that displays a gradual shift in cellular and extracellular properties from the tendon side through to the bone side [1-5]. Yet, despite its importance, the formation of this cellular gradient as well as the underlying molecular mechanism remain largely unknown.

In recent years, the initial events that lead to the formation of the embryonic attachment unit (AU), which serves as the primordium of the enthesis, have started to be investigated. These studies identified the progenitors of the AU and showed that they express both the chondrogenic and tenogenic transcription factors *Sox9* and scleraxis (*Scx*), respectively [6, 7]. The patterning of the *Sox9*^*+*^*/Scx*^*+*^ progenitors along the skeleton is regulated by a genetic program that includes several transcription factors [8]. Later, the *Sox9*^*+*^*/Scx*^*+*^ cells differentiate into *Gli1*^*+*^ cells [9-11]. Furthermore, both molecular and mechanical signals regulate the AU. TGFβ signaling regulates the specification of AU progenitors, whereas BMP and FGF signaling as well as mechanical signals determine their fate and differentiation [7, 12, 13]. Postnatal enthesis cells have been termed fibrocartilage cells based on their histological appearance, since they display morphological features that are shared with tenocytes and chondrocytes [14]. In recent years, several studies have identified some of the genes that these cells express, including collagens type I, II and X, Indian hedgehog (*Ihh*), parathyroid hormone-related peptide (*PTHrP*), patched 1 (*Ptc1*), runt related transcription factor 2 (*Runx2*), tenascin C (*Tnc*), and biglycan (*Bgn*) [14-17]. Interestingly, these genes are also expressed by cells in the neighboring tissues, namely by chondrocytes or tenocytes. However, despite these advances, a comprehensive molecular signature of this tissue and the mechanism that enables its formation are still missing. In this work, we aimed to decipher the identity of the fibrocartilage cells that form the attachment tissue between tendon and bone. Transcriptomic analysis of the attachment cells, which was validated by *in situ* hybridization (ISH) and single-molecule fluorescent ISH (smFISH), showed that these cells express a mix of the transcriptomes of chondrocytes and tenocytes. Chromatin analysis further verified the transcriptomic results and provided a mechanistic explanation for the bi-fated behavior of attachment cells, which share enhancers with their neighboring tenocytes or chondrocytes. Finally, we identify the transcription factors KLF2 and KLF4 as regulators of attachment cell differentiation. Overall, we provide the transcriptional as well as the epigenetic mechanism that allows attachment cells to activate a combination of cartilage and tendon transcriptomes and, thereby, the formation of the unique transitional tissue.

## Results

### Attachment cell transcriptome is a mix of chondrocyte and tenocyte transcriptomes

To date, the transcriptome of attachment cells has not been characterized thoroughly. We therefore analyzed the transcriptome of embryonic day (E) 14.5 attachment cells from the prominent deltoid tuberosity and greater tuberosity of the humerus (Fig. S1A,B). With the goal to isolate these cells specifically, we generated a compound mouse by crossing three mouse lines, namely *Col2a1-Cre, R26R-tdTomato* and *Scx-GFP* (see Materials and Methods) *[6, 7]*. Thus, the fluorescent reporter tdTomato labeled *Col2a1*-expressing chondrocytes, whereas GFP fluorescently labeled *Scx*-expressing tenocytes. Unexpectedly, the two reporters failed to label the attachment cells that were located in between these two populations. This failure might be due to a missing regulatory element in one of the constructs that were used to produce each transgenic reporter. Nevertheless, the borders between tendon and attachment cells and between cartilage and attachment cells were clearly demarcated. We therefore used laser capture microdissection (LCM) to subdivide the attachment site into three cellular compartments, namely attachment cells, adjacent tenocytes and adjacent chondrocytes. As controls, samples were also taken from two more compartments, remote tenocytes and remote chondrocytes.

Initial analysis of the different transcriptomes using principal components analysis (PCA) showed that the transcriptomes of tenocytes and chondrocytes were clearly separated, whereas attachment cells were located between the two cell types, recapitulating their anatomical positions. This suggests that the attachment cell transcriptome is largely shared with both chondrocytes and tenocytes (Fig. 1A, PC1 52.47%).

**Figure 1:**
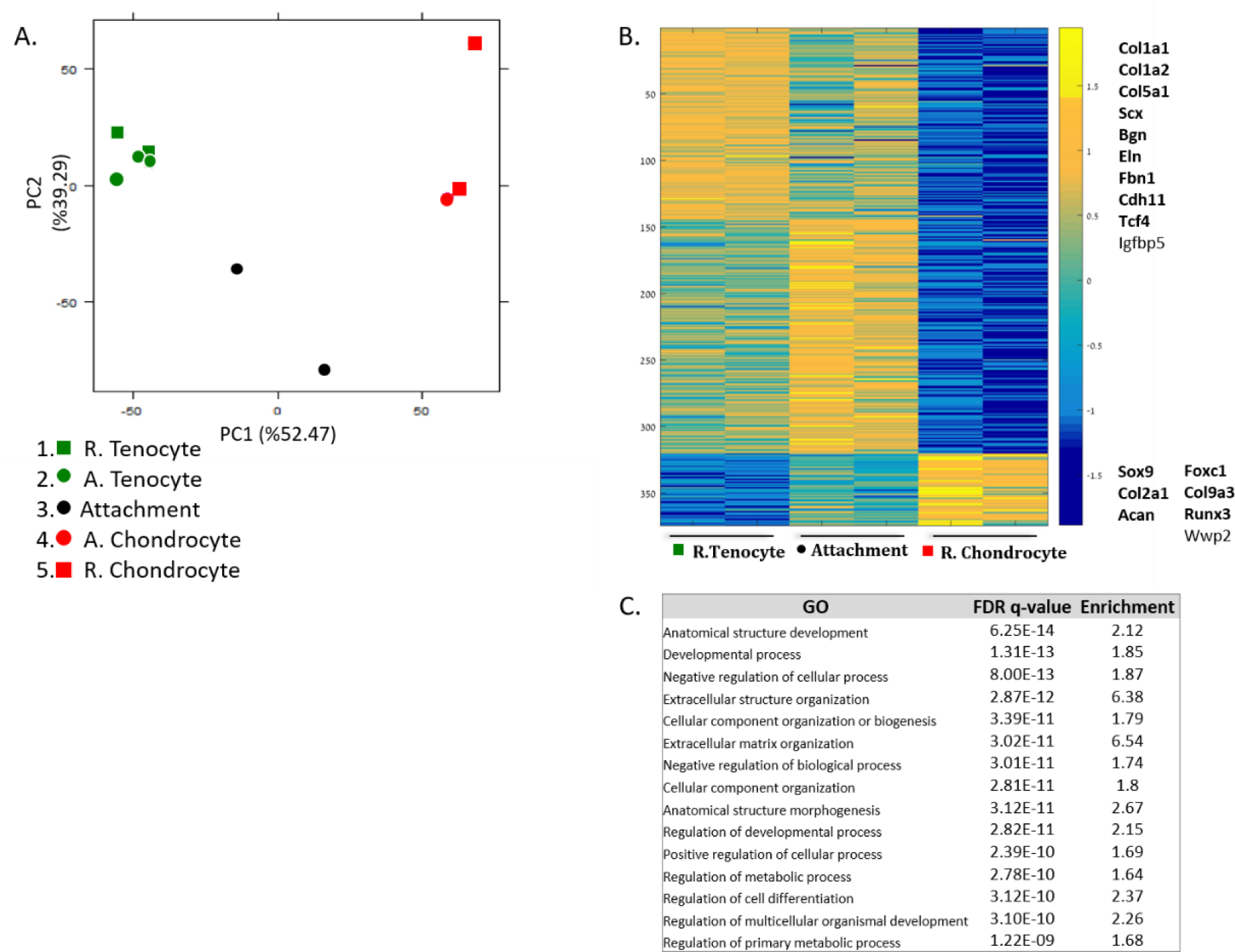
Transcriptomic analysis of tendon-to-bone attachment site domains at E14.5. **A.** Principal component analysis (PCA) of MARS-Seq data from E14.5 attachment site samples. The *x*-axis (PC1) shows the highest variance among the samples. Interestingly, the samples are arranged according to their anatomical locations. “R” (samples 1 and 5) stands for remote and “A” (samples 2 and 4) is for adjacent. The *y*-axis (PC2) shows that tenocytes and chondrocytes are closer to one another, while attachment cells (black circle) were found to be remote from both of them, i.e. with higher variance, suggesting a unique gene expression profile. **B.** Heatmap of gene expression profiles at E14.5 shows 374 selected genes that exhibited differential expression between tenocytes and chondrocytes and were also expressed by attachment cells. Color bar (−1.5-0-1.5) represents the log-normalized counts standardized per gene, as yellow is higher than the mean (0) and blue is lower than the mean. Attachment cells display a gradient of gene expression profiles, reflecting their function as a transitional tissue. The upper cluster contains genes highly expressed in tenocytes (e.g. *Co1a1, Col1a2, Col5a1, Scx, Bgn*), whereas the lower cluster contains genes highly expressed in chondrocytes (e.g. *Sox9, Col2a1, Acan*). Top list on the right contains genes found to be expressed in attachment cells and in tenocytes, whereas bottom list contains genes expressed in attachment cells and chondrocytes; genes in bold type are known tenocyte or chondrocyte markers. **C.** List of the top15 GO terms (biological process) associated with the 374 shared genes.

To further support our initial observation that the transcriptome of the attachment cells is a mixture of chondrocyte and tenocyte transcriptomes, we clustered the statistically significant differentially expressed genes between all samples into 5 clusters, using CLICK (Fig. S3 and see Materials and Methods). Out of 865 identified genes, 735 genes were found in two clusters. The first cluster contained mainly known tenogenic genes and the second contained chondrogenic genes (Fig. S2). From these two clusters, 374 genes, 320 of them tenogenic and 54 chondrogenic, were also found to be expressed by attachment cells. They included major regulators and marker genes of the two tissues, such as *Sox9, Col2a1* and *Acan* for chondrocytes and *Col1a1, Col1a2, Scx* and *Col5a1* for tenocytes (Fig. 1B). GO analysis of these shared genes yielded terms relating to anatomical structure development, developmental process, negative regulation of cellular process, extracellular structure organization and others (Fig. 1C).

The third cluster comprised 54 genes that were found to be up-regulated in cartilage adjacent to attachment cells alone (Fig. S2). Interestingly, our analysis identified 24 and 23 genes that were found to be down- or up-regulated in attachment cells, shown by the fourth and fifth clusters, respectively (Fig. S2). The genes that were found to be uniquely up-regulated in attachment cells included transcription factors, such as the Krüppel-like factors (KLFs), *Lmo1* and *Gli1*, which could act as regulators of the genetic program of attachment cells. In addition, this set included differentiation markers such as *Thy1*, regulators of bone e.g. *Acp5* and *Alpl*, protein kinases such as *Mapk12* and *Mast2*, and signaling molecules such as *Nod, Traip, Aplnr* and others (Fig. S3A,B). GO analysis of these genes yielded terms relating to regulation of cytokine and IL-12 production, as well as response to laminar fluid shear stress (Fig. S3C). These results clearly show that the transcriptome of the attachment cells includes a mixture of tenocyte and chondrocyte genes, many of which are involved in ECM organization and developmental processes, in addition to a unique subgroup of genes that are up-regulated in these cells.

### Attachment cells co-express tenocyte and chondrocyte genes

To validate our transcriptome analysis, we performed single- and double-fluorescent *in situ* hybridization (FISH) using marker genes for tenocytes and chondrocytes that were selected from the transcriptomic results, namely *Igfbp5*, biglycan (*Bgn*), *Col5a1* and *Col1a1* for tenocytes, and *Col11a1* and *Wwp2* for chondrocytes. As seen in Figures 2 and S4, in agreement with the transcriptome analysis, the selected markers were co-expressed by the attachment cells. To further substantiate this result, we performed single-molecule FISH to study *in vivo* expression of *Wwp2* (chondrogenic marker) and *Bgn* (tenogenic marker) at a single-cell resolution. Results showed that attachment cells co-expressed both markers at the single-cell level (Fig. 2M,N). Overall, these results support the transcriptome analysis, indicating that the attachment cell population expresses in parallel both chondrogenic and tenogenic genes in the same cell to form the attachment site (referred to in the following as mixed transcriptome or mixed gene expression).

**Figure 2:**
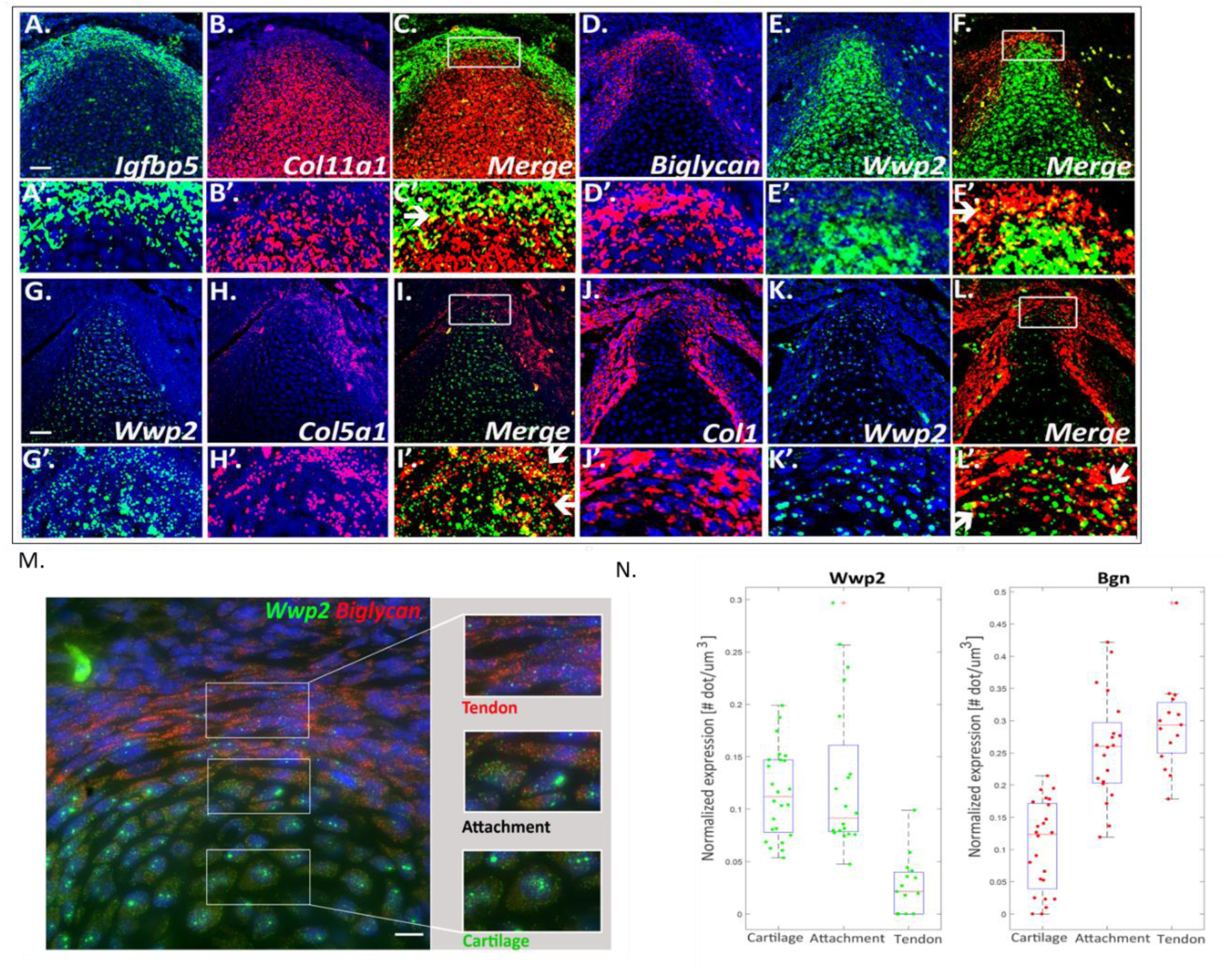
Attachment cells co-express tendon and cartilage genes at the single-cell level. **A-L.** Double-fluorescent ISH for mRNA of tendon (*Igfbp5*, biglycan, *Col5a1, Col1a1*) and cartilage (*Col11a1* or *Wwp2*) genes shows that attachment cells (in yellow, shown by arrows) exhibit an *in vivo* gene expression profile that combines tendon and cartilage genetic programs. A-L: X20 magnification, 50 µm scale bar; A’-L’: magnification of upper panels. **M.** Single-molecule fluorescent ISH (smFISH) of mRNA of tendon biglycan (*Bgn*, red) and cartilage *Wwp2* (green) genes on the background of DAPI staining (blue) further validates the dFISH results. X100 magnification, 10 µm scale bar. **N.** Quantification of *Bgn* and *Wwp2* smFISH results in cartilage, attachment site and tendon.

### Genome-wide profiling of attachment cell-specific regulatory regions

To gain a mechanistic understanding of how attachment cells activate a combination of two transcriptomes, we compared chromatin accessibility in these cells with open chromatin signatures defining chondrocytes and tenocytes by conducting an assay for transposase-accessible chromatin with high-throughput sequencing (ATAC-Seq) [18]. This method allows to profile open chromatin regions, some of which may act as enhancers. To isolate E13.5 humeral deltoid tuberosity tenocytes and attachment cells, we generated another compound mouse line harboring *Sox9-CreER, tdTomato* and *Scx-GFP* transgenes (Fig. S1C-F). Additionally, chondrocytes were FACS-sorted from E13.5 *Col2a1-CreER*^*T*^*-tdTomato-Scx-GFP* mouse. These three cell populations were then subjected to ATAC-Seq (see Materials and Methods).

Initial PCA analysis of accessible chromatin profiles for each FACS-sorted cell population once again revealed that tenocytes and chondrocytes were clearly separated, while attachment cells resided between these two cell types (Fig. S5A, Fig. 1A). Next, we compared global chromatin accessibility among the three cell types by calculating the level of overlap among the ATAC-Seq peaks (Fig. 3A). While the majority of the peaks were shared by all three cell types, attachment cells had a significantly lower number of unique peaks (*p* < 1e-4 relative to both cell types, chi-square with Yates correction), and a significantly higher overlap with the other two cell types (*p* < 2.2e-1).

**Figure 3:**
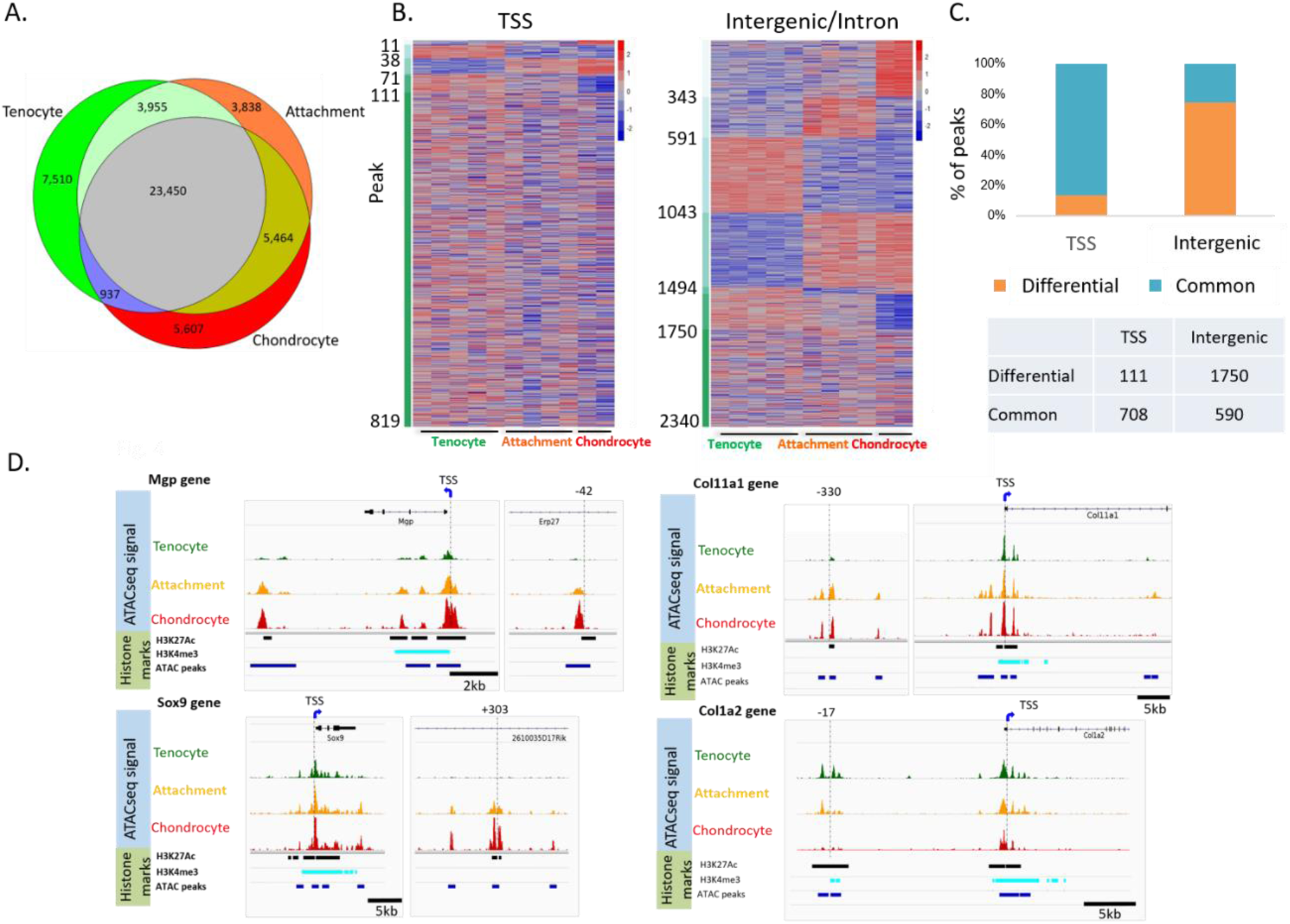
Accessible chromatin reveals an epigenetic mechanism shared by attachment cells and neighboring tenocytes or chondrocytes. **A.** Venn diagram showing cell-specific or overlapping peaks of ATAC-Seq among tenocytes, chondrocytes and attachment cells. **B.** Heatmap of ATAC-Seq peaks associated with E14.5 differentially expressed genes. Left: TSS peaks, right: intergenic or intron peaks. The peaks are sorted according to their degree of accessibility across the three cell types. **C.** Percentage of common peaks (shared by three cell types) vs. differential peaks (the chromatin is open only in one or two cell types) compared between TSS and intergenic areas (P=0, chi-square test). **D.** IGV snapshots of the TSS region of *Mgp, Sox9, Col11a1* and *Col1a2* genes, as well as potential enhancers of these genes.

Analysis of the ATAC-Seq signal revealed that 13,017 peaks were located near transcription start sites (TSSs), whereas 31,856 peaks were in intergenic or intron regions. Most of the peaks that were located near TSSs were accessible in all three cell types (87%), and only 13% were accessible in one or two cell types (Fig. S5C,D). Next, we studied the ATAC-Seq signal of peaks associated with the genes that were differentially expressed at E14.5, using HOMER default parameters. We found that 819 peaks were located near transcription start sites (TSSs), whereas 2340 peaks were in intergenic or intron regions. Most of the peaks that were located near TSSs (708, 86%) were accessible in all three cell types, and only 111 (13%) were accessible in one or two cell types (Fig. 3B,C). This low level of differential accessibility is inconsistent with the possibility that promoter accessibility is the main mechanism regulating the bi-fated attachment cells. Interestingly, a significantly higher fraction of intergenic peaks were specific to one or two cell types (1750, 74.7%, *p* = 0, chi-square test, Fig. 3B,C). Moreover, ∼46% of the intergenic peaks that were differentially accessible between tenocytes and chondrocytes were also found to be accessible in attachment cells. Overall, these results suggest that the intergenic elements that are shared between attachment cells and chondrocytes or tenocytes may act as enhancers that regulate the mixed transcriptome of attachment cells.

To identify such dual cell type-specific enhancers likely regulating attachment cell differentiation, we next screened for shared enhancers of 15 *bona fide* markers of tenocytes or chondrocytes that were found to be expressed in E14.5 attachment cells (Fig. 1B). To improve the prediction of these enhancers, we selected our ATAC-Seq peaks based on their proximity to genes with verified expression in attachment cells and another cell type (Fig. S4, Fig. 2) and computationally intersected them with ENCODE datasets of histone modification marks associated with enhancers and promoters in mouse limbs at E13.5 (H3K27Ac, H3K9ac, H3K4me3, H3K4me1 ChIP-Seq), and other datasets [19, 20] (Fig. 4 and Table 1), revealing the degree of evolutionary conservation of each core sequence [21]. For example, as shown in Figure 3D, we identified a region at −42 kb from the TSS of Mgp, a *bona fide* chondrogenic marker [22], which was accessible in chondrocytes and attachment cells, whereas in tenocytes this site was closed. Another example is *Sox9*, a *bona fide* chondrogenic marker. At +303 kb from *Sox9*, we identified a region that was accessible in attachment cells and chondrocytes, but not in tenocytes. The same pattern was observed for a region at −330 kb from the TSS of a third *bona fide* chondrogenic marker, namely *Col11a1* [23]. The opposite pattern was observed at −17 kb from the TSS of *Col1a2*, a *bona fide* tenogenic marker, where we identified a region that was accessible in attachment cells and tenocytes, but not in chondrocytes. Similar results were obtained for additional chondrogenic markers, such as *Sox6*, and for tenogenic markers *Tnc* and *Col1a1* (data not shown). Importantly, we found that the chromatin accessibility patterns of these putative enhancers were in agreement with the transcriptomic and ISH results, as shown, for example, by *Sox9* and *Col5a1* (Figs. 1B, 2, S4). This suggests that the mechanism for the activation of a mixed transcriptome in attachment cells is based on sharing regulatory elements with chondrocytes and tenocytes.

**Table 1:** Summary of information used to predict enhancer activity *in vivo*. Transgenic mouse transgenic reporter assay (LacZ) of elements that were screened at E14.5 forelimbs. The following criteria were considered to predict enhancers that may be active at E14.5 (top to bottom): Distance from TSS, coordinate, conservation and verification by H3K27Ac, H3K4me1 ChiP-seq of ENCODE, HiC results (Andrey et al., 2017) and core sequence conservation for the following eight elements (left to right): Col1a1 element, Klf2 element, Sox9 element, Mgp element, Col11a1 element, two negative elements in the forelimb for Eln gene, in addition to Igfbp5 element (the last with verified activity in the nose area). We first compared the ATAC-seq data to MARS-seq transcriptome analysis and chose peaks that were assigned to genes with differential expression (using GREAT). According to the ATAC-seq results, the TSS of most of these genes was accessible in all three tissue types. Therefore, we searched for putative enhancers that could regulate the differential gene expression. We then compared the elements overlap with E13.5-E14.5 histone marks (H3K27Ac or H3K4me1 of ENCODE) and HiC results (Andrey et al., 2017), to increase the probability of *in vivo* verification, in addition to preliminary results of gene expression (i.e. the chosen elements were assigned to genes of interest, which were identified by RNA-seq and validated by ISH). The degree of evolutionary conservation of the core sequence was also taken into consideration while prioritizing the elements.

|  | mEL4,<br>Col1a1 | mEL5,<br>Klf2 | mEL6,<br>Sox9 | mEL7,<br>Mgp | mEL8,<br>Col11a1 | mEL10,<br>mEL12, Eln | mEL3,<br>Igfbp5 |
| --- | --- | --- | --- | --- | --- | --- | --- |
| **Distance** | -63,010 | -30,343 | 303,066 | -42,288 | -331,264 | -3,433<br>-43,487 | -3,605 |
| **Coordinate** | Chr11:94872064 -<br>94874364 | Chr8: 72287698-72291754 | Chr11:113084290-113086290 | Chr6:136917343- 136918843 | Chr3: 113698476-113700076 | Chr5: 134749924-134751424<br>Chr5: 134790228-134791228 | Chr1: 72877526-72879452 |
| **Encode**<br>H3K27ac/H3K4me1 | + | + | + | + | + | + | + |
| **HiC** |  | + |  |  |  |  |  |
| **Core seq**<br>**conservation** | opossum | chicken | platypus | opossum | lizard | opossum | chicken |
| **Klf2 binding**<br>**site** | + | + | + | - | - |  |  |

**Figure 4:**
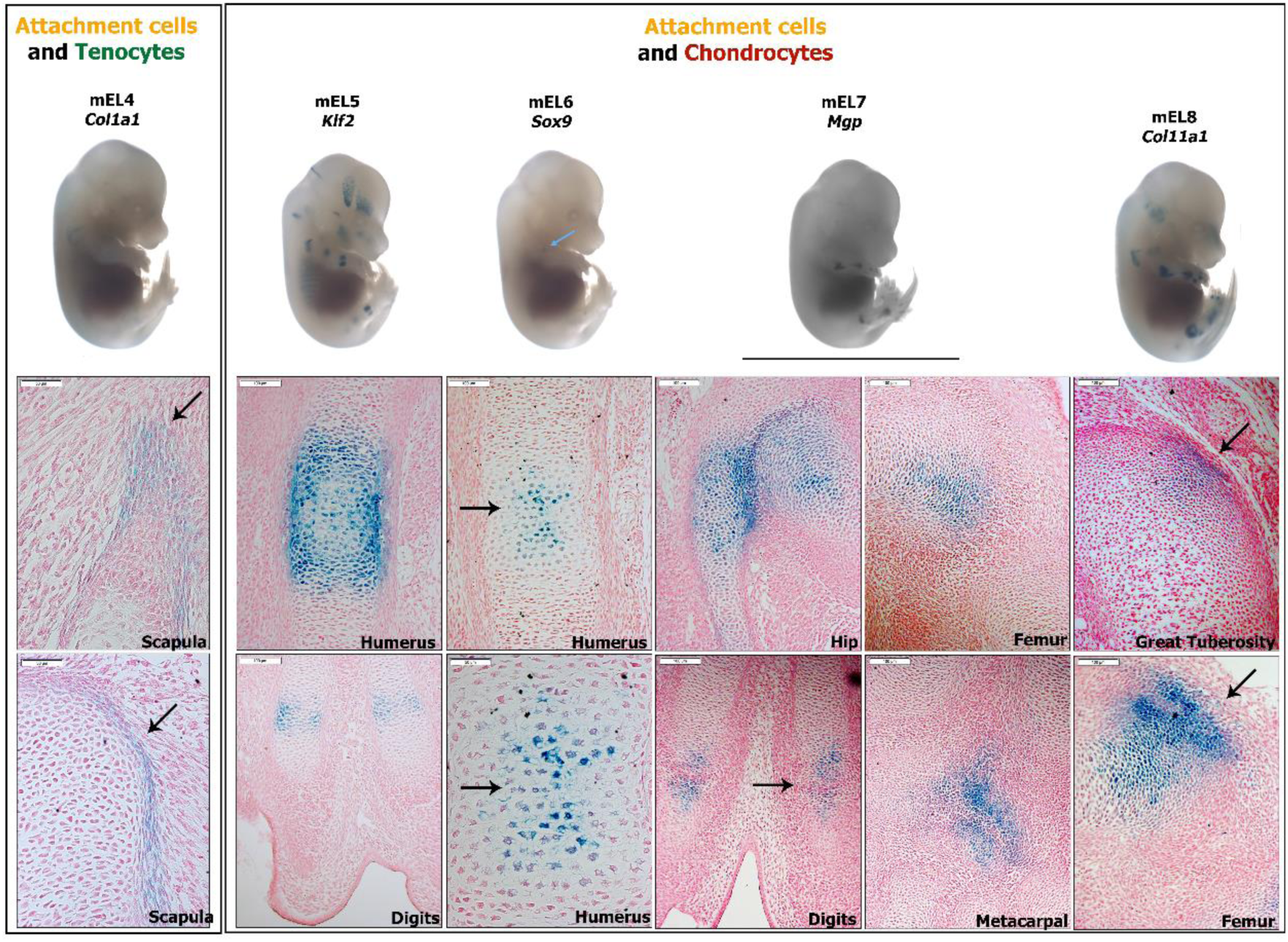
*In vivo* analysis of enhancers identifies shared domains of activity between attachment cells and neighboring tissues. Transgenic mouse reporter enhancer assay (lacZ) of elements positive at E14.5 (marked in light blue; for each enhancer, an E14.5 embryo and forelimb and/or hindlimb sagittal sections are shown). Left to right: *Col1a1* element activity is seen at the teres major insertion at the scapula. *Klf2* element activity is seen in hypertrophic chondrocytes and perichondrium at the humerus and forelimb digits. *Sox9* element activity is seen in hypertrophic chondrocytes of the humerus. *Mgp* element activity is seen in the hip, digit and metacarpals joints in addition to the posterior distal side of the femur. *Col11a1* element activity is seen in the greater tuberosity insertion and anterior distal side of the femur.

### Shared regulatory elements drive expression in attachment cells and flanking cartilage or tendon cells

The identification of multiple predicted enhancer regions near genes expressed by attachment cells and tenocytes or chondrocytes suggests that attachment cells are regulated predominantly by enhancers with shared activities. To test this hypothesis *in vivo*, we took advantage of a recently developed, site-directed transgenic mouse enhancer reporter system (Kvon et al. submitted). Using this system, we studied the activity of eight elements, which were selected because they were associated with *bona fide* marker genes for tenocytes or chondrocytes, and were found to be expressed in the attachment cells. Moreover, they were predicted to drive transcription in attachment cells and one of the flanking tissues (Fig. 3D). The activity of these representative elements was examined at E14.5, a stage at which chondrocytes, tenocytes, and attachment cells have already been established.

Five elements were found to be active in the mouse forelimbs, as well as in other anatomical areas (Fig. 4). The *Col1a1-*associated element drove *lacZ* expression at the teres major insertion into the scapula, in agreement with the ATAC-Seq results, which predicted its activity in tenocytes and attachment cells. This result suggests that *Col1a1* element is active in tenocytes and attachment cells. *Klf2* and *Sox9* elements were predicted to be active in chondrocytes and attachment cells (Fig. 3D). The *Klf2* element indeed drove reporter activity in hypertrophic chondrocytes and perichondrium at the humerus and forelimb digits, in addition to the skull and mandible (Fig.4). The *Sox9* element showed activity solely in hypertrophic chondrocytes of the humerus. These results suggest a chondrocyte-specific function of these two enhancers. *Mgp* element was also predicted to be active in chondrocytes and attachment cells (Fig. 3D). Its activity was seen in forelimb and hindlimb, specifically in hip, metacarpals joints and digits as well as in the posterior distal side of the femur, a site where ligaments (e.g. the cruciate ligaments) are inserted into the femur at the knee area and at ligament insertion into to the hip (e.g. iliofemoral ligament) (Fig. 4), verifying its activity in chondrocytes and attachment cells. Lastly, *Col11a1* element activity was predicted in chondrocytes and attachment cells (Fig. 3D). Its activity verified the bioinformatic analysis, showing LacZ staining in the greater tuberosity insertion, as well as in the posterior side of the skull, the nasal bone area and the anterior distal side of the femur (Fig. 4). These results therefore suggest *Mgp* and *Col11a1* elements are active in chondrocytes and attachment cells, in addition to *Col1a1* element which is active in tenocytes and attachment cells, as the chromatin analysis predicts. Overall, these results provide a proof of concept for the ability of the accessible intergenic elements we have identified to act as enhancers that drive expression in both attachment cells and chondrocytes or tenocytes. This supports our hypothesis that shared enhancers activate a mixed transcriptome in attachment cells.

### Krüppel-like factors are regulators of attachment cell development

Our finding of enhancers that can drive the transcription of the mixed transcriptome of the attachment cells raised the question of the identity of the transcription factors (TFs) that can potentially bind to these elements. To identify such factors, we used Genomatix to analyze accessible elements that were associated with differentially expressed genes for over-representation of transcription factor binding sites (TFBS), selecting the top 50 TFBS families, and then mining our transcriptomic data for the expression of these TFBS families (see Materials and Methods). Among the differentially expressed genes at E14.5 we identified NFIs (*Nfia*), GLIs (*Gli1*), KLFs (*Klf2* and *Klf4)*, ZBTBs (*Zbtb48*) and RUNXs (*Runx3*, Table 2, Fig. 5A), whose expression was up-regulated in attachment cells. Further support for these results was provided by HOMER motif analysis [24], which showed significant over-representation of KLFs and RUNXs TFBSs. We therefore sought to explore the possible role of KLFs as regulators of attachment cells.

**Table 2.**
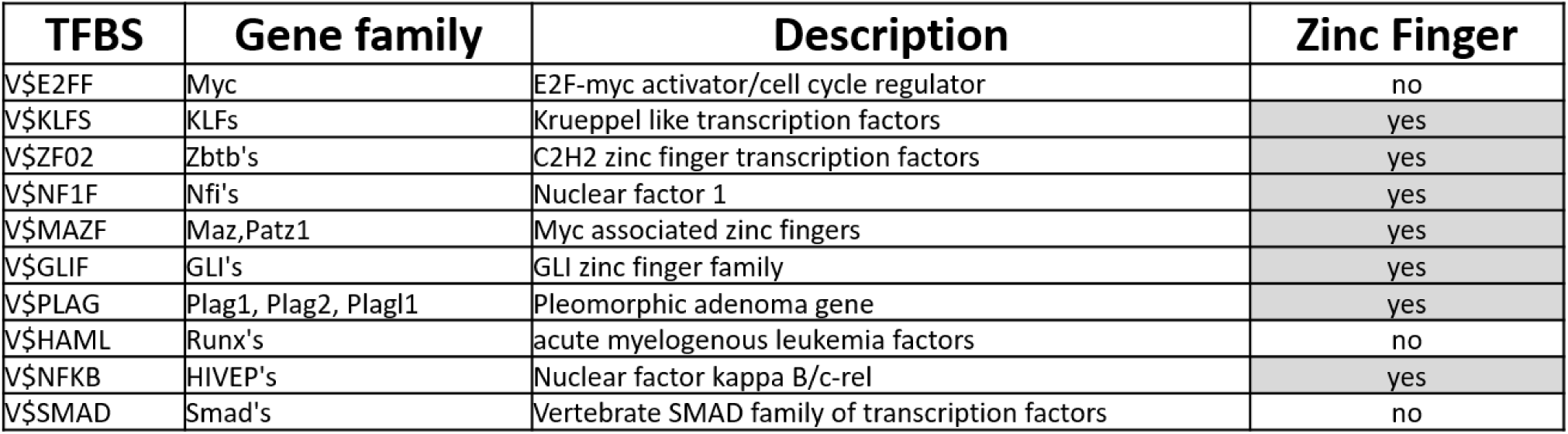
Genomatix analysis of the genomic regions of cis-regulatory elements, identified by ATAC-Seq. Over-representation of transcription factor binding sites (TFBS) families were identified. Crossing these results with E14.5 transcriptome revealed differentially expressed TFs from the KLFs (*Klf4* and *Klf2*), GLIs (*Gli11*), NFIs (*Nfia*), ZBTBs (*Zbtb48*) and RUNXs families.

| TFBS | Gene family | Description | Zinc Finger |
| --- | --- | --- | --- |
| V\$E2FF | Myc | E2F-myc activator/cell cycle regulator | no |
| V\$KLFS | KLFs | Krueppel like transcription factors | yes |
| V\$ZF02 | Zbtb's | C2H2 zinc finger transcription factors | yes |
| V\$NF1F | Nfi's | Nuclear factor 1 | yes |
| V\$MAZF | Maz,Patz1 | Myc associated zinc fingers | yes |
| V\$GLIF | GLI's | GLI zinc finger family | yes |
| V\$PLAG | Plag1, Plag2, Plagl1 | Pleomorphic adenoma gene | yes |
| V\$HAML | Runx's | acute myelogenous leukemia factors | no |
| V\$NFKB | HIVeP's | Nuclear factor kappa B/c-rel | yes |
| V\$SMAD | Smad's | Vertebrate SMAD family of transcription factors | no |

**Figure 5:**
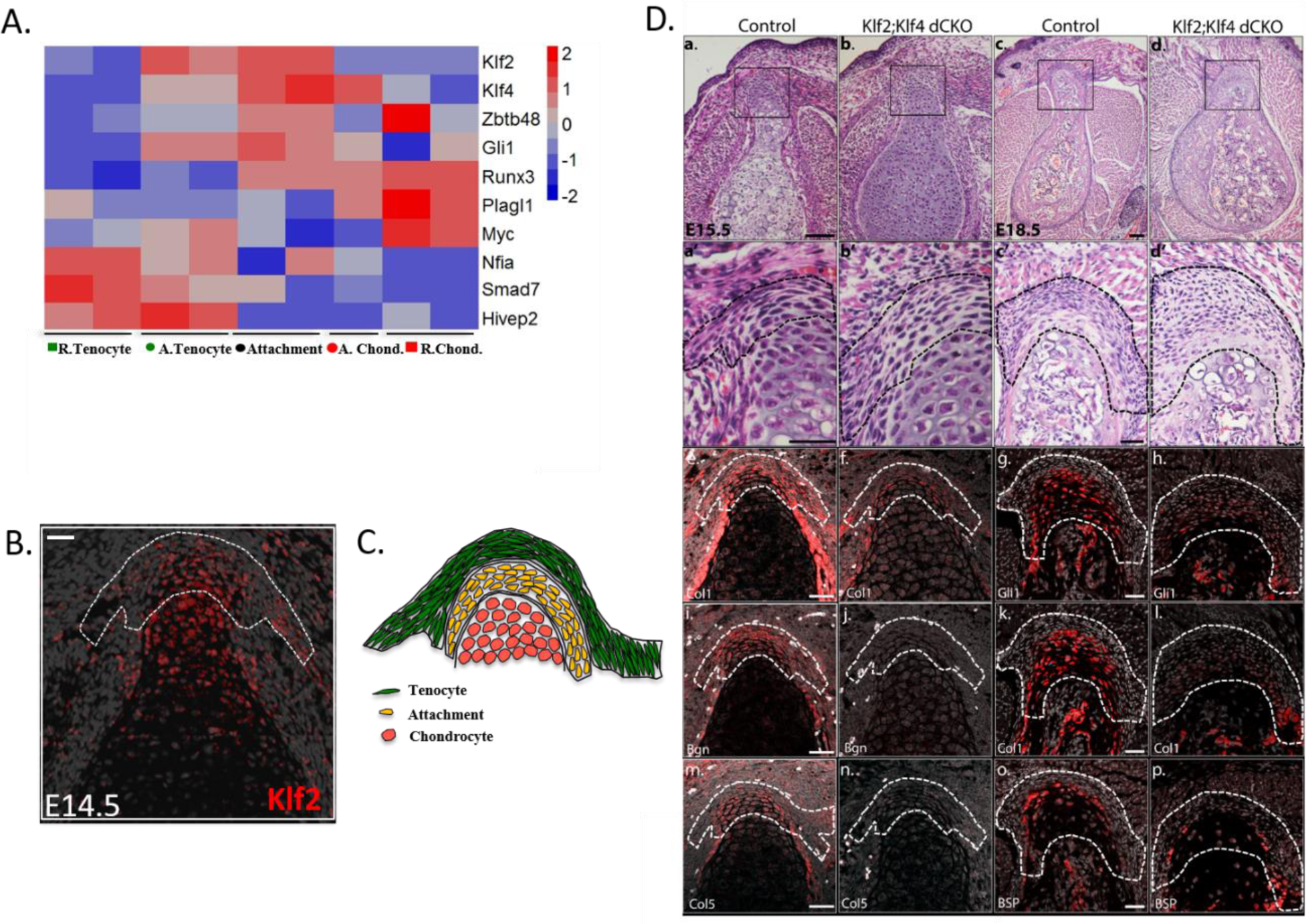
Krüppel-like factors (KLFs) are regulators of attachment cell development. **A.** Heatmap of selected transcription factors at E14.5. Transcriptome analysis shows up-regulated expression *of Klf2, Klf4* and *Gli1* in attachment cells. **B.** E14.5 ISH validated these results, showing *Klf2* expression in attachment cells (X20 magnification, 25 µm scale bar). **C.** Scheme of attachment site. **D.** KLF2 and KLF4 are regulators of attachment cell development. a-d. Histological transverse sections through the humeral deltoid tuberosity of E15.5 *Prx1-Klf2-Klf4* and E18.5 *Prx1-Klf2-Klf4* mutant and control embryos (X10 and X5 magnification, 100 µm scale bar). a’-d’. Higher magnification of upper panel (X40 for E15.5 and X20 for E18.5, 40 µm scale bar). e-p. ISH for *Col1a1* and *Bgn* genes of E15.5 *Prx1-Klf2-Klf4* mutant and control embryos. ISH for *Gli1, Col1a1* and *Bsp* genes of E18.5 *Prx1-Klf2-Klf4* mutant and control embryos (X20 magnification, 40 µm scale bar).

Focusing on *Klf2*, which was found to be differentially expressed specifically in the attachment site, we first validated its expression in the forming attachment site by *in situ* hybridization (Fig. 5B). Next, we analyzed the enhancers that were shown by the bioinformatic analysis to be active in attachment cells and either tenocytes or chondrocytes (Fig. 4). Three of these enhancers had Klf2 binding sites in their sequence (Table 1), further supporting a potential role for KLFs during attachment site development.

Previous studies demonstrated that Klf2 and Klf4 are functionally redundant, as KLF4 has ∼90% sequence similarity to KLF2 in its zinc finger DNA binding domain, suggesting that these factors could have common target sequences [25]. We therefore proceeded to study attachment cell development upon blocking the expression of both *Klf2* and *Klf4* in limb mesenchyme, using *Prx1-Cre* as a deleter and focusing on E15.5 and E18.5, a period during which the attachment site of the deltoid tuberosity undergoes differentiation and consequently grows in size. Transverse histological sections through the deltoid tuberosity of E15.5 control mice showed that the attachment cells were packed together and surrounded by ECM (Fig. 5Da’). In contrast, in the putative attachment site of *Prx1-Klf2-Klf4* double conditional knockout (dcKO) embryos, the cells were sparse with reduced ECM (Fig. 5Db’). By E18.5, this difference was more pronounced (Fig. 5Dc’,d’). To gain a molecular understanding, we studied the expression of several genes that were previously shown to be expressed at these stages in the attachment site [11] (Fig. 2). Indeed, we found that the expression of *Col1a1, Gli1, Bsp, Bgn* and *Col5a1* was reduced in the dcKO attachment site, relative to the control (Fig. 5De-p), supporting a role for KLF2/4 in the attachment site.

Finally, to further validate the involvement of KLFs in activation of gene expression in the attachment site, we searched for KLF2/4 binding sites in ATAC-Seq peaks associated with the 374 genes that were shown to be expressed by attachment cells (Fig. 1C). Interestingly, we found that many of these genes had KLF2/4 binding sites in their regulatory regions (72% of the 374 attachment genes relative to 53% in the whole genome, *p* < 1e-4, chi-square). We then searched for KLF2/4 binding sites in ATAC-Seq peaks that were associated with genes whose expression was reduced in the dcKO attachment site (Fig.5De-p). For *Gli1*, we found KLF2/4 TFBSs in peaks that reside −2.1 and −1.5 kb from its TSS (Table 3). For *Col5a1*, we found multiple binding sites for KLF2 or KLF4. Together, these results indicate that KLF2/4 play an essential role in regulating attachment cell gene expression.

**Table 3.**
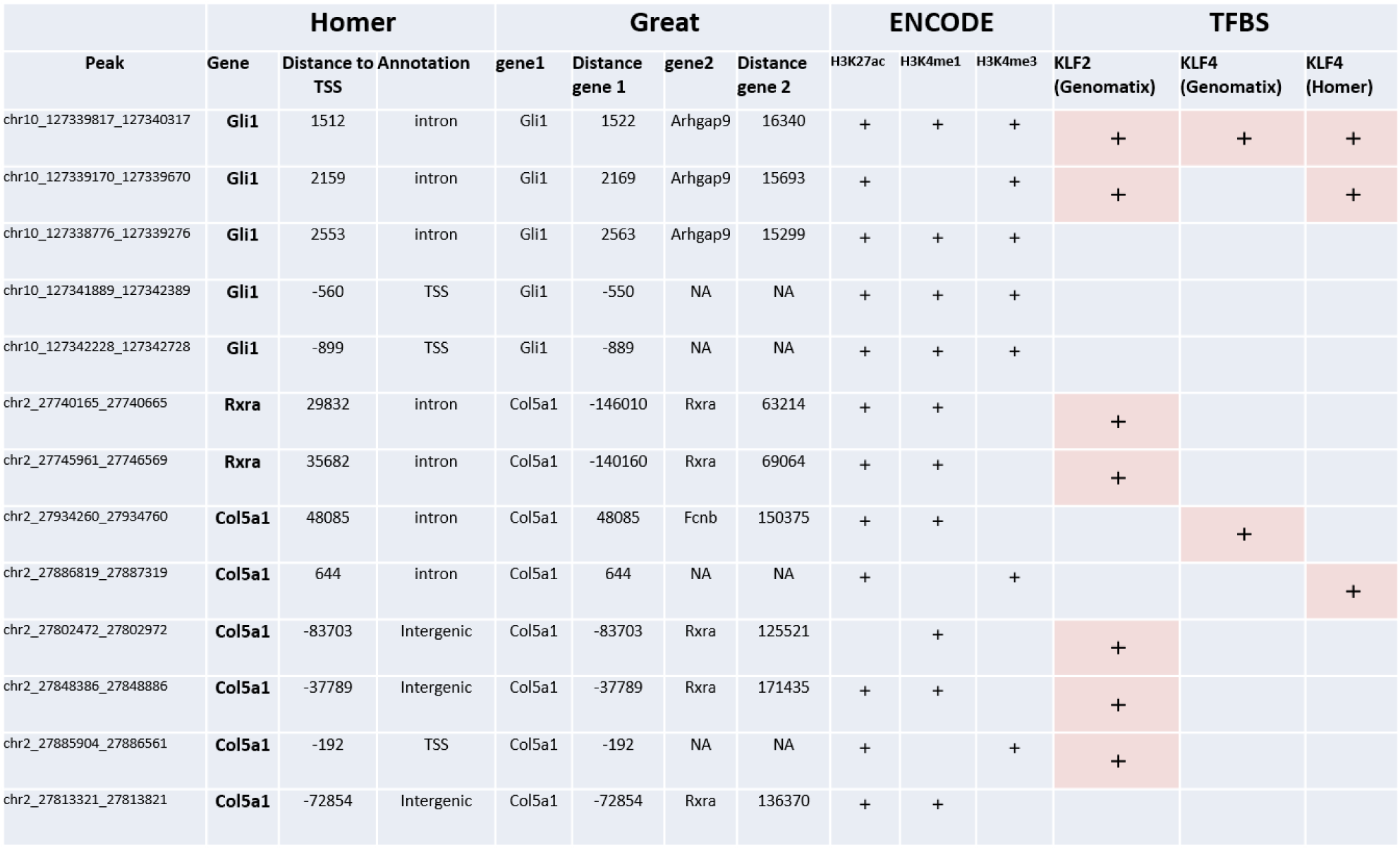
Gli1 and Col5a1 genes are expressed in the attachment site and have TFBS for KLFs. Search for KLF2/4 binding sites in ATAC-Seq peaks that were associated with genes whose expression was reduced in the dcKO attachment site (Fig.5De-p). For *Gli1*, we found KLF2/4 TFBSs in peaks that reside −2.1 and −1.5 kb from its TSS (indicated as + under TFBS, pink). For *Col5a1*, we found multiple binding sites for KLF2 or KLF4 (indicated as + under TFBS, pink).

Combined with the bioinformatic analysis of chromatin and transcriptomic data, these results suggest that KLF2/4 are major regulators of tendon-to-bone attachment, playing a central role in attachment cell differentiation.

## Discussion

In this work, we describe the unique transcriptome that allows cells of the attachment between tendon and bone to act as a transitional tissue. The ability to activate a combination of chondrogenic and tenogenic transcriptomes is regulated by sharing enhancers with these cells. Finally, we identify the transcription factors KLF2/4 as regulators of these unique bi-fated cells.

The existence of borders between tissues that differ in cell type, extracellular matrix composition, structure and function raises the question of how tissues are connected. The border can be sharp, as seen in blood vessels, where pericytes and endothelial cells are separated by a basement membrane, or between the esophagus and the stomach in the gastrointestinal tract [26, 27]. On the other hand, the border can be less defined histologically and molecularly, thus forming a transitional tissue. Examples for the latter are the borders between the sections of the small intestine and between tendon and bone [27-29]. From a broader perspective, as all organs and systems are made of different tissues, understanding the biology of border tissues is imperative. Moreover, some of these border tissues are involved in various pathologies. For example, gastric cancers may emerge from distinct anatomical areas, such as the esophagus-stomach boundary [27, 30, 31]. Another congenital disease affecting this boundary tissue is anophthalmia-esophageal-genital (AEG) syndrome, which results from *SOX2* loss-of-function mutation, causing esophageal atresia, i.e. esophagus obstruction [32]. This involvement further underscores the need to study border tissues.

In the case of the attachment between tendon and bone, the significance of this tissue is demonstrated by enthesopathies, a collective name for injuries and pathologies of the enthesis. For example, over 30% of the population over the age of 60 will injure their shoulder’s rotator cuff [33]. Failure rates of surgical reattachment range from 20% for small tears to 94% for repair of massive tears [34, 35]. The high failure and recurrence rates of these procedures highlight the need for understanding the biology of this complex transitional tissue of the enthesis. This understanding may allow the development of new strategies to improve the treatment of enthesopathies.

There are two options to form a transitional tissue. The first strategy is by mixing cells from the two neighboring tissues, such as in the epithelia of the urinary tract [28]. Alternatively, the border cells can express a mixture of the transcriptomes of the two neighboring cell types. As we show here, the attachment cells represent the latter strategy well, as they express a high number of genes that are differentially expressed by either tenocytes or chondrocytes. These cells display morphological features that are shared with tenocytes and chondrocytes [14]. Our results therefore provide a molecular explanation for the age-old histological definition of enthesis cells as fibrocartilage, which was based on their morphology [14]. Moreover, the finding of mixed matrix genes in the transcriptome of the attachment cells may provide a mechanism for the formation of a transitional tissue, which allows safe transfer of forces by the tendon between muscle and bone. Interestingly, the attachment cells express more tenogenic than chondrogenic genes, suggesting that they may be part of the connective tissue lineage. In addition to expression of chondrogenic and tenogenic genes, we identified genes that are uniquely expressed by attachment cells. These genes may provide another level of specificity to the regulation of the development of this unique tissue.

Our finding that attachment cells are bi-fated raises the question of the mechanism that underlies this fate. An immediate implication of our finding is that there must be an epigenetic mechanism that supports the bi-fated state. The observed chromatin accessibility at the sites of the promoters of most of the shared genes in all three cell types rules out the possibility of limited promoter accessibility as the main mechanism. By contrast, the high percentage of shared intergenic sites between attachment cells and one group of flanking cells, i.e. chondrocytes or tenocytes, suggests that this is the main mechanism. Moreover, many of these shared sites correlated with ENCODE datasets, where they appear as putative enhancers. Finally, we found three different enhancers that can drive gene expression in attachment cells and either tendon or cartilage. Overall, these findings strongly support our hypothesis that the regulatory mechanism is based on the ability of attachment cells to share enhancers with either chondrocytes or tenocytes in order to drive the mixed expression profile of these bi-fated cells.

Sharing enhancers is not the only possible strategy for the generation of mixed transcriptome. A simple alternative would be a specific set of enhancers to be used by the attachment cells. A possible explanation for the sharing strategy is the common origin of all these cells, which is limb mesenchyme originating from lateral plate mesoderm [36, 37]. It is possible that during development, limb mesenchymal progenitors display highly accessible chromatin; yet, during differentiation, this accessibility is restricted to prevent the expression of genes from alternate lineages. In contrast to this restriction process, in the bi-fated attachment cells the shared sites are maintained accessible to allow the expression of the mixed transcriptome. A mechanism for silencing of genes of alternate lineages was previously described. For example, polycomb-repressed chromatin leads to silencing of genes of alternate lineages, leading to the commitment to a specific cell fate [38-40]. Interestingly, previous studies demonstrated the importance of chromatin repression in the developing limb, showing how deletion of *Ezh2*, which acts as the enzymatically active subunit of PRC2, leads to skeletal malformations [41]. This obviously raises the question of the mechanism that prevents this silencing in attachment cells.

It is clear that we cannot exclude the possibility that an active mechanism, such as the SWI/SNF remodeling complexes, opens the chromatin structure in bi-fated cells to allow attachment cell dual behavior [42]. However, such a mechanism cannot explain why the strategy of shared enhancers was selected. Overall, our results reveal a novel function for chromatin state, which allows the activation of two sets of genes in a third cell type to create a new cell fate that forms a transitional tissue.

KLF2 and KLF4 are known to regulate several biological processes, such as promoting the differentiation of gut and skin (KLF4, [43, 44] as well as the immune system (KLF2, [45]), maintaining pluripotency of embryonic stem cells (KLF2 and KLF 4, [46]), and, together with other factors, inducing pluripotency to generate iPSC by reprogramming (KLF4, [47]). Several works describe the involvement of KLF2 and KLF4 in the musculoskeletal system. In bones, *Klf4* over-expression in osteoblasts caused delayed bone development, in addition to impaired blood vessel invasion and osteoclast recruitment [48]. Another study showed that KLF2/4 are expressed during chick limb development in tendons and ligaments as part of the genetic program that regulates connective tissues [49].

Previous studies showed that KLF2 and KLF4 display high similarity in protein sequences [50, 51], suggesting that these factors could have common target sequences and may be functionally redundant. Indeed, loss of both KLF2 and KLF4 during embryogenesis led to abnormal blood vessel development and early lethality. This phenotype was more severe than what was observed in embryos that lost only KLF2 or KLF4 [25]. Furthermore, Orgeur et al. (2018) identified 313 target genes shared between KLF2 and KLf4, suggesting that they overlap in regulating gene expression.

In this work, we show that KLF2/4 are central regulators of the attachment site. While the attachment did form initially in their absence, the subsequent differentiation failed, suggesting that KLF2/4 play a role at this stage. While we concentrated in this study on the attachment site, it is most likely that KLF2/4 play a role also in other musculoskeletal tissues such as the skeleton, tendon and muscle. This possibility is supported by previous studies, where KLF2/4 were shown to be expressed in osteoblasts, chondrocytes, tenocytes and muscle connective tissues [48, 49].

Previous studies demonstrate the role of muscle-induced mechanical load in the development of attachment site [13]. In that context, our finding that KLF2/4 regulate the differentiation of attachment cells is interesting, since previous works have shown that these factors are mechanically regulated. It was shown in mice that shear stress on the vessels induced by blood flow leads to up-regulation of *Klf2* expression [52]. Additional *in vitro* studies showed that KLF2 and KLF4 are influenced by shear stress [25, 53, 54]. It is therefore possible that these factors are regulated by muscle forces, leading to the proper differentiation and maturation of the attachment site.

The ability of KLF2/4 to regulate gene expression in the attachment site is supported by our finding that many of the genes that were expressed by attachment cells had in their regulatory region KLF2/4 binding sites. Yet, it is clear that not all of them share this property, suggesting that these two factors are part of a larger transcriptional network. For example, our bioinformatic analysis identified other TF families such as GLI’s, RUNX’s and NFI’s as regulators of the attachment sites. *Gli1* was previously reported as a marker for enthesis cells [9, 11, 17, 55]. Since gene expression by attachment cells is regulated by sharing enhancers with chondrocytes or tenocytes, it is reasonable to assume that some regulators of these cells might be part of the network that regulates the attachment cells. Indeed, loss of the tendon regulator *Scx* in mice led to failure of attachment cells to differentiate. Additionally, loss of the chondrogenic regulator *Sox9* in *Scx*-expressing cells led to failure in attachment site formation [7, 13].

To conclude, by characterizing the transcriptome and chromatin landscape of tendon-to-bone attachment cells, we provide a molecular understanding of the bi-fated identity of these cells. Moreover, by identifying the transcription factors KLF2/4 as central regulators and the strategy of sharing enhancers with either tenocytes or chondrocytes, we provide a mechanism that regulates these bi-fated cells (Fig. 6). These findings present a new concept for the formation of a border tissue, which is based on the simultaneous expression of a mixed transcriptome of the two flanking cell types by the intermediate cells. This strategy allows the formation of a unique transitional tissue without developing *de novo* a dedicated genetic program that regulates a third, new cell fate.

**Figure 6:**
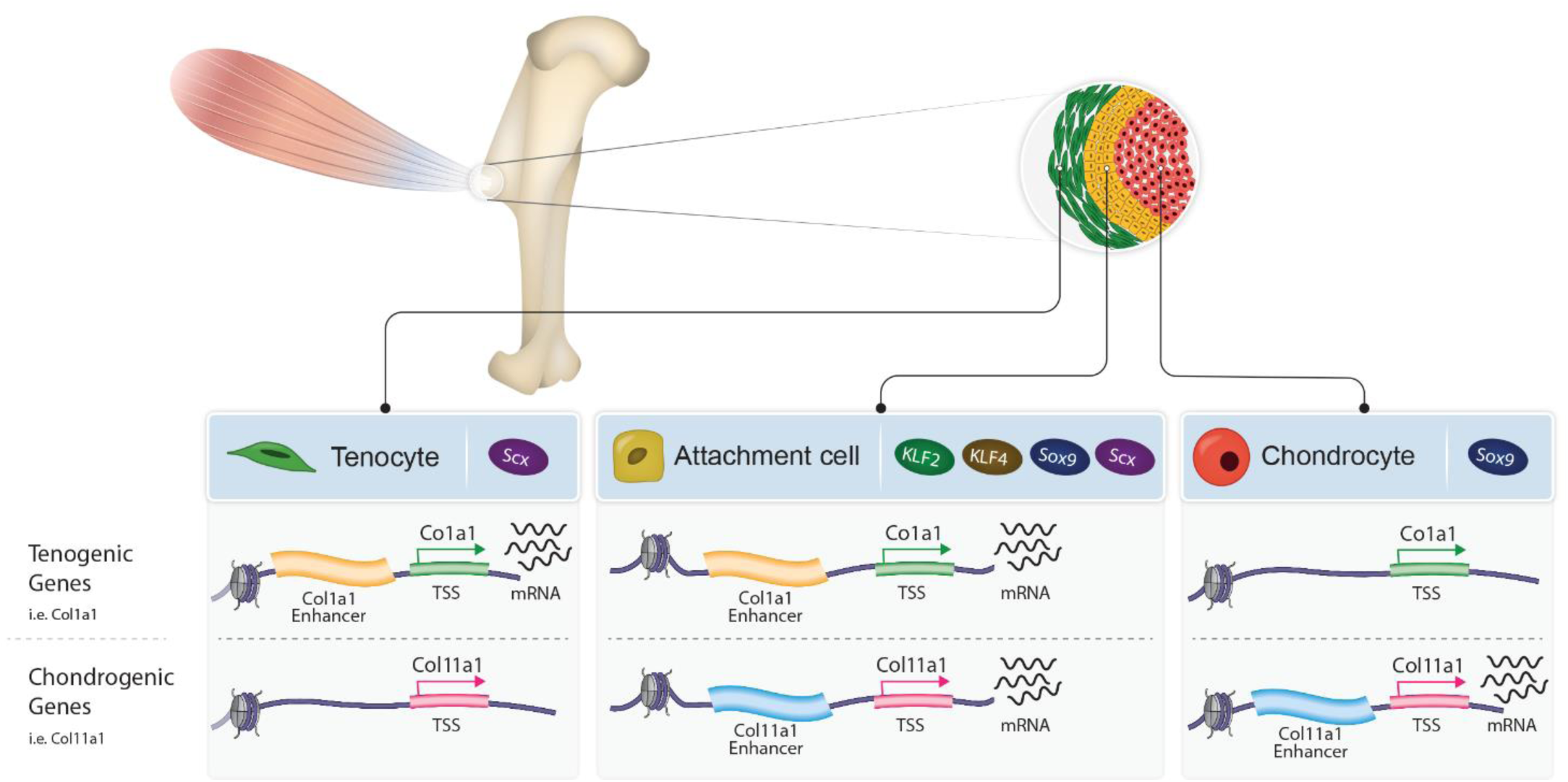
Proposed model of bi-fated tendon-to-bone attachment cells are regulated by shared enhancers and KLF transcription factors. Tenocytes (green, left) and chondrocytes (red, right) express tenogenic (i.e. *Col1a1*) or chondrogenic (i.e. *Col11a1*) genes, respectively, whereas attachment cells (yellow, middle) express both chondrogenic and tenogenic genes to form the attachment site. Attachment cells duality of gene expression is regulated epigenetically by intergenic chromatin areas, which are accessible in these cells and in either tenocytes or chondrocytes. Additionally, at the transcriptional level, the transcription factors KLF2/4 are expressed by attachment cells and regulate their differentiation.

## Materials and Methods

### Ethics declaration

All mice were maintained and used in accordance with protocols approved by the Weizmann Institutional Animal Care and Use Committee (IACUC).

All animal work was reviewed and approved by the Lawrence Berkeley National Laboratory (LBNL) Animal Welfare Committee. All mice used in this study were housed at the Animal Care Facility (ACF) at LBNL. Mice were monitored daily for food and water intake, and animals were inspected weekly by the Chair of the Animal Welfare and Research Committee and the head of the animal facility in consultation with the veterinary staff. The LBNL ACF is accredited by the American Association for the Accreditation of Laboratory Animal Care (AAALAC). Transgenic mouse assays were performed in *Mus musculus* FVB background mice.

### Animals

The generation of floxed *Klf2 [52]*, floxed *Klf4 [44], Prx1-Cre [56] Sox9-CreER [57], Col2-CreER*^*T*^ *[58], Col2a1-Cre [59], R26R-tdTomato [60]* and *Scx-GFP [61]* mice have been described previously.

To create *Col2a1-CreER, R26R-tdTomato, Sox9-CreER, R26R-tdTomato* and *Col2a1-Cre, R26R-tdTomato* reporter mice, all on the background of *Scx-GFP*, floxed *R26R-tdTomato* mice were mated with *Col2a1-CreER, Sox9-CreER* or *Col2a1-Cre* mice, respectively. These strains were mated on a mixed background of C57BL/6 and B6.129 (ICR) mice and used for LCM and FACS experiments. To create *Prx1-Klf2-Klf4* mutant mice, floxed *Klf2-Klf4* mice were mated with *Prx1-Klf2-Klf4* mice. As a control, we used embryos that lack *Cre* alleles. E14.5 wild-type C57BL/6 mice were used for *in situ* hybridization experiments as well.

For FACS experiments, *Sox9-CreER* or *Col2a1-CreER* mice were crossed with *Rosa26-tdTomato* reporter mice, all on the background of *Scx-GFP*. Induction of Cre recombinase was performed at various pregnancy stages by administration of 0.03 mg/gr tamoxifen/body weight in corn oil by oral gavage (stock concentration was 5 mg/ml). In all timed pregnancies, plug date was defined as E0.5. For harvesting of embryos, timed-pregnant females were sacrificed by cervical dislocation. Tail genomic DNA was used for genotyping by PCR.

### Laser capture microdissection (LCM)

E14.5 *Col2a1-tdTomato-Scx-GFP* mouse forelimbs were dissected, shortly fixated in 4% PFA, washed with PBS and cryo-embedded (as described by Bhattacherjee et al., 2004). Next, samples were cryo-sectioned and mounted on LCM slides (PET, Zeiss), washed with RNase-free water and EtOH (Arcturus dehydration component kit) according to an altered protocol of Pazin et al [62]. LCM (PALM MicroBeam C system, Zeiss) was calibrated for refined tissue cutting. Isolated cells were collected to LCM caps (Adhesive Cap 500 clear, Zeiss) and RNA was purified using RNeasy FFPE Kit (Qiagen).

### Fluorescence-activated cell sorting (FACS)

Flow cytometry analysis and sorting were performed at the Weizmann Institute of Science Flow Cytometry Core Facility on a BD FACS AriaIII instrument (BD Immunocytometry Systems) equipped with 488, 407, 561 and 633 nm lasers, using a 70 μm nozzle, controlled by BD FACS Diva software v8.0.1 (BD Biosciences). Further analysis was performed using FlowJo software v10.2 (Tree Star). For collection of cells, *Sox9-CreER*^*T2*^*-tdTomato;ScxGFP* or *Col2a1-CreER-tdTomato;ScxGFP* mice were crossed with *Rosa26-tdTomato;ScxGFP* reporter mice. Embryos were harvested at E13.5 following tamoxifen administration at E12.0, as described above. Forelimbs were dissected and suspended in cold PBS. To extract cells from tissues, PBS was replaced with 1 ml heated 0.05% trypsin and collagenase type V (dissolved in DMEM, Sigma) and incubated for 15 min at 37°C, gently agitated every 5 minutes. Tissues were then dissociated by vigorous pipetting using 1 ml tips. Next, 4 ml of DMEM supplemented with 10% FBS and 1% Pen-Strep was added and cell suspensions were filtered with 40 μm filter net. Finally, tubes were centrifuged at 1,000 rpm for 7 minutes, supernatant was removed and cells were resuspended in 1 ml of cold PBS and used immediately for FACS. Single-stained GFP and tdTomato control cells were used for configuration and determining gate boundaries. Live cells were gated by size and granularity using FSC-A versus SSC-A and according to DAPI staining (1 μg/ml). FSC-W versus FSC-A was used to further distinguish single cells. In addition, unstained, GFP-stained only and tdTomato-stained only cells were mixed in various combinations to verify that the analysis excluded false-positive doublets. GFP was detected by excitation at 488 nm and collection of emission using 502 longpass (LP) and 530/30 bandpass (BP) filters. tdTomato was detected by excitation at 561 nm and collection of emission using a 582/15 BP filter. DAPI was detected by excitation at 407 nm and collection of emission using a 450/40 BP filter.

### Real-time PCR (RT-PCR)

Total RNA was purified from LCM-isolated samples of E14.5 mouse forelimbs using RNeasy FFPE Kit (Qiagen). Reverse transcription was performed with High Capacity Reverse Transcription Kit (Applied Biosystems) according to the manufacturer’s protocol. Analysis of *Col2a1* and *Scx* was performed to monitor RNA quality during LCM calibrations, whereas RNA quantity was monitored by analysis of β–actin. RT-PCR was performed using Fast SYBR Green master mix (Applied Biosystems) on the StepOnePlus machine (Applied Biosystems). Values were calculated using the StepOne software. Data were normalized to *18S* rRNA or β-actin in all cases.

### *In situ* hybridization

Section ISH were performed as described previously [63]. Single-and double-fluorescent ISH on paraffin sections were performed using DIG- and/or FITC-labeled probes [64]. After hybridization, slides were washed, quenched and blocked. Probes were detected by incubation with anti-DIG-POD (Roche; 1:300) and anti-FITC-POD (Roche, 1:200), followed by Cy2-tyramide- and Cy3-tyramide-labeled fluorescent dyes according to the instructions of the TSA Plus Fluorescent Systems Kit (Perkin Elmer).

### Single-molecule fluorescent *in situ* hybridization (smFISH)

Harvested E14.5 forelimbs were fixed with cold 4% formaldehyde (FA) in PBS and incubated first in 4% FA/PBS for 3 h, then in 30% sucrose in 4% FA/PBS overnight at 4°C with constant agitation. Fixed tissues were embedded in OCT and sectioned at a thickness of 10 µm. The preparation of the probe library, hybridization procedure and imaging conditions were previously described [65-67]. In brief, probe libraries were designed against biglycan (*Bgn*) and *Wwp2* mRNA sequences using the Stellaris FISH Probe Designer (Biosearch Technologies, Inc., Petaluma, CA) coupled to Quasar 670 and CAL Fluor Red 610, respectively. Libraries consisted of 17-96 probes each of length 20 bps, complementary to the coding sequence of each gene (Table S4). Nuclei were stained with DAPI. To detect cell borders, Alexa Fluor 488 conjugated phalloidin (Thermo Fisher, A12379) was added to the GLOX buffer, which was wash for 15 minutes. Slides were mounted using ProLong Gold (Molecular Probes, P36934).

### Image acquisition and analysis

For smFISH image acquisition, we used a Nikon-Ti-E inverted fluorescence microscope equipped with a Photometrics Pixis 1024 CCD camera to image 10-µm-thick cryosections. For image analysis we used ImageM, a custom MATLAB program [66], which was used to compute single-cell mRNA concentrations by segmenting each cell manually according to the cell borders and the nucleus. The size of the nucleus was detected automatically by the program according to the DAPI signal. For each cell, the concentration of cytoplasmic mRNA of each gene was calculated by measuring the number of dots per volume. Images were visualized and processed using ImageJ 1.51h [68] and Adobe Illustrator CC2018.

### RNA sequencing

For this analysis, we performed a bulk adaptation of the MARS-Seq protocol [69] [70] to generate RNA-seq libraries for expression profiling of the purified RNA from E14.5 LCM-isolated samples. Briefly, RNA from each sample was barcoded during reverse transcription and pooled. Following Agencourct Ampure XP beads cleanup (Beckman Coulter), the pooled samples underwent second strand synthesis and were linearly amplified by T7 *in vitro* transcription. The resulting RNA was fragmented and converted into a sequencing-ready library by tagging the samples with Illumina sequences during ligation, RT, and PCR. Libraries were quantified by Qubit and TapeStation as well as by qPCR for actb gene as previously described [69, 70]. Sequencing was done on a Hiseq 2500 SR50 cycles kit (Illumina).

The data were analyzed using the Pipeline Pilot-designed pipeline for transSeq (by INCPM,https://incpmpm.atlassian.net/wiki/spaces/PUB/pages/36405284/tranSeq+on+Pip eline-Pilot). Briefly, the analysis included adapter trimming, mapping to the mm9 genome, collapsing of reads with the same unique molecular identifiers (UMI) of 4 bases (R2) and counting of the number of reads per gene with HTseq-count [71], using the most 3’ 1000 bp of each RefSeq’s transcript. DESeq2 [72] was used for differential expression analysis with betaPrior set to true, cooksCutoff=FALSE, independentFiltering=FALSE. Benjamini-Hochberg method was used to adjust the raw p-values for multiple testing. Genes with adjusted p-value ≤ 0.05 and fold change ≥ 2 between every two conditions were considered as differential. Clustering of the normalized read count of differentially expressed genes was done using click algorithm (Expander package, [73]), followed by visualization by R (R Core Team, 2013). Further analysis was performed using GSEA (Broad institute) and Gorilla [74, 75].

### Assay for transposase-accessible chromatin with high-throughput sequencing

ATAC-Seq data were trimmed from their adaptors and filtered from low quality reads using Cutadapt followed by alignment to the mm10 genome (GRCm38.p5) using Bowtie2 (version 2.3.4.1) [76]. PCR-duplicate reads were removed with Picard ‘MarkDuplicates’ (http://broadinstitute.github.io/picard/). Mitochondrial reads were removed from the alignment, and the data were further filtered to contain only reads with a unique mapping with SAMtools (-F 4 -f 0×2). Read pairs with inner distance of up to 120 bp were selected as representing the accessible chromatin region. MACS2 (version 2.1.1.20160309) [77] was applied for peak calling using the setting: callpeak -f BAMPE--nomodel. Peaks from all samples were combined and merged with BEDTools [78], followed by extension to a minimum length of 500 bp. For every tissue, a set of reproducible peaks was obtained by voting, which means that a normalized read count ≥ 30 was detected in at least 50% of the replicates. Peaks that were not reproducible in any tissue were removed. Peaks that reside in the ENCODE “Blacklist” regions, i.e. regions that were previously found by ENCODE (PMID: 22955616) to produce artificial signal (http://mitra.stanford.edu/kundaje/akundaje/release/blacklists/), were also eliminated. Peak quantification was done with BedTools [78] following by DESeq2 [72] normalization. Peaks with an averaged normalized read count ≥ 30 in at least one of the studied tissues were selected for the downstream analyses.

The crude data of the work has been deposited on NCBI GEO (GSE144306).

#### Annotation and genomic feature enrichment analysis

Annotation of ATAC-Seq peaks was performed using HOMER [24] and GREAT [79]. When a peak was associated by GREAT to multiple genes, the two closest genes were selected for further analysis. ATAC-Seq peaks that were at a distance of up to −2 kb down or +0.5 kb up from a TSS of their annotated gene (HOMER) were considered as promoter peaks; otherwise, peaks were considered as distal. To rank distal ATAC-Seq peaks as putative cis-regulatory elements, we calculated the overlap between the peaks and relevant histone modification datasets (ChIP-seq) performed by the ENCODE project [80] on E13.5 C57BL/6 mouse embryo limb. The overlap was calculated using BEDTools intersect [78]. The following datasets were used: ENCSR905FFU (H3K27ac) and ENCSR426EZM (H3K4me1) as markers of enhancers, ENCSR416OYH (H3K4me3) as a marker of promoters and ENCSR022DED (H3K9me3). Overlap with the phastConsElements60wayPlacental track downloaded from the UCSC site [21] was calculated to account for evolutionary conservation. Enrichment analysis of over-representation of TFBSs in the ATAC-Seq peaks was performed with the RegionMiner tool of Genomatix™ and HOMER.

### Enhancer reporter assays in mouse embryos

Candidate enhancers were PCR-amplified and cloned upstream of a *Shh*-promoter-LacZ-reporter cassette. We used a mouse enhancer-reporter assay that relies on site-specific integration of a transgene into the mouse genome [Kvon et al 2020, submitted, 82]. In this assay, the reporter cassette is flanked by homology arms targeting the H11 safe harbor locus [81]. Cas9 protein and a sgRNA targeting H11 were co-injected into the pronucleus of FVB single cell-stage mouse embryos (E0.5) together with the reporter vector [Kvon et al 2020, submitted, 82]. Embryos were sampled and stained at E14.5. Embryos were only excluded from further analysis if they did not carry the reporter transgene. All mouse work was reviewed and approved by the Lawrence Berkeley National Laboratory Animal Welfare and Research Committee.

## Acknowledgments

We thank Nitzan Konstantin for expert editorial assistance, Dr. Douglas Lutz and service engineer Tal Alon (Getter Bio-Med, Zeiss) for LCM calibration, Neria Sharabi from the Department of Veterinary Resources, Weizmann Institute, and all Zelzer lab members for suggestions and advice. We thank E. Sebzda for providing floxed *Klf2* mice and the Mutant Mouse Regional Resource Center (MMRRC) at UC Davis for providing floxed *Klf4* mice. We thank the ENCODE Consortium and the ENCODE production laboratory for generating the described datasets.

This study was supported by grants from the National Institutes of Health (R01 AR055580), the David and Fela Shapell Family Center for Genetic Disorders, the David and Fela Shapell Family Foundation INCPM Fund for Preclinical Studies, and the Estate of Bernard Bishin for the WIS-Clalit Program (to E.Z.). Work at Lawrence Berkeley National Lab was supported by National Institute of Health grant R01HG003988 (A.V.) and performed under Department of Energy Contract DE-AC02-05CH11231, University of California.

## Supplementary

**Figure S1:**
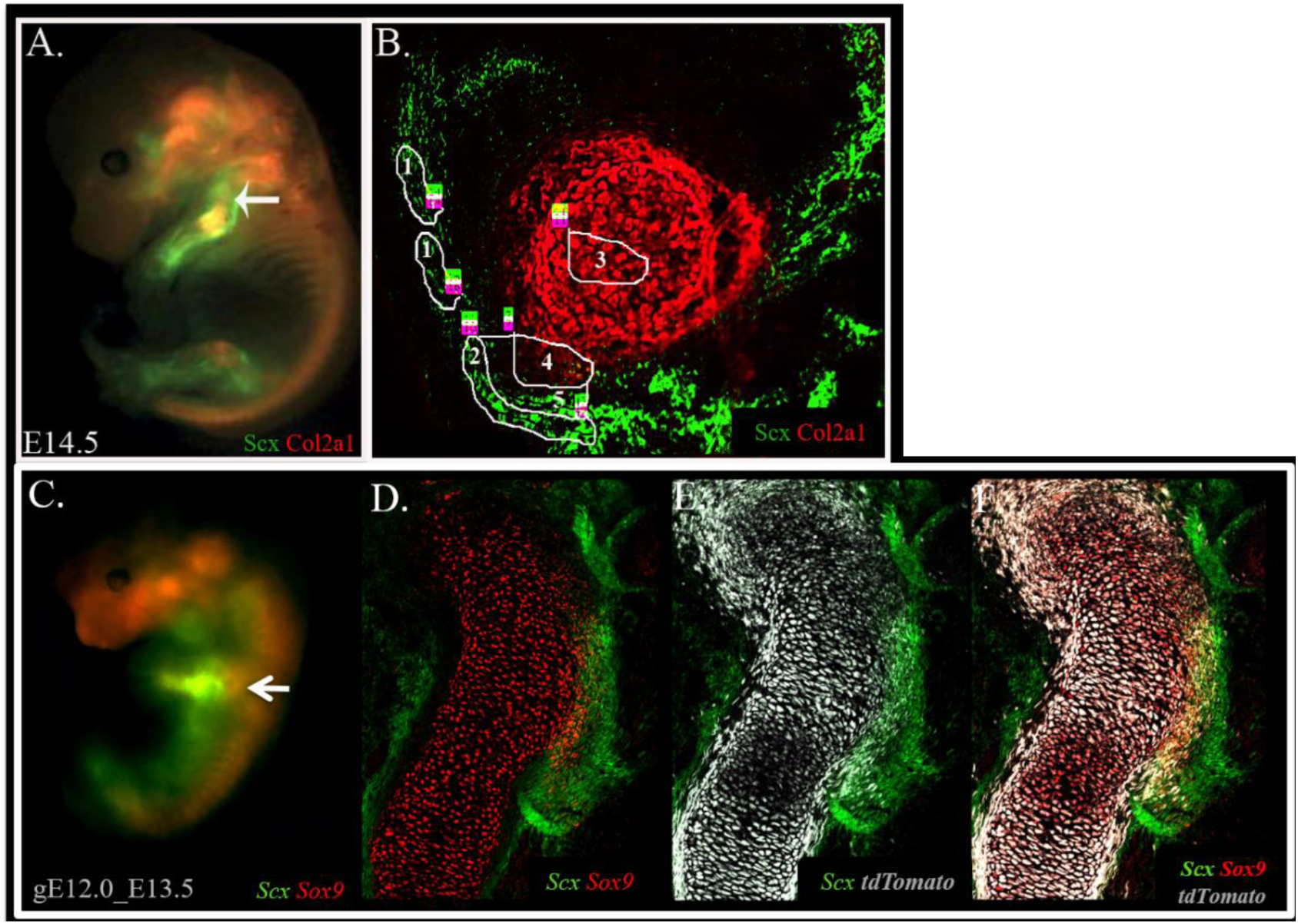
Analysis of the embryonic tendon-to-bone attachment site in a triple-transgenic mouse line. **(A)** The attachment site at the prominent deltoid tuberosity and greater tuberosity (arrow) of the humerus was analyzed in E14.5 Col2a1-Cre-tdTomato-Scx-GFP transgenic mice. The fluorescent reporter tdTomato labeled *Col2a1*-expressing chondrocytes, GFP fluorescently labeled Scx-expressing tenocytes. Unexpectedly, the two reporters failed to label the attachment cells that were located in between these two populations. Nevertheless, the borders between tendon and attachment cells and between cartilage and attachment cells were clearly demarcated. **(B)** Cells from five distinct areas in and around the tendon-to-bone attachment site were isolated for RNA sequencing using LCM: 1, remote tenocytes; 2, adjacent tenocytes; 3, remote chondrocytes; 4, adjacent chondrocytes; 5, attachment cells. The attachment site was divided into three cellular compartments as follows: adj. chondrocytes, defined as the bone eminence protruding from the primary cartilaginous element of the humerus and marked by Col2a1; tendon, defined as the side adjacent to tendon cells and marked by Scx-GFP; and attachment cells, which were located between the two other compartments and were negative to both reporters. Two additional groups of cells were isolated by LCM as a control, namely cells from tendon and cartilage tissues that were distant from the attachment site, referred to in the following as remote tenocyte and remote chondrocyte (part of the primary cartilaginous element of the humerus), respectively. **(C-F)** Attachment cells at the humeral deltoid tuberosity and greater tuberosity of E13.5 Sox9-CreER-tdTomato-Scx-GFP embryos were isolated by FACS for ATAC-seq analysis. The cells connecting tendon to bone are Scx-Sox9 double-positive.

**Figure S2:**
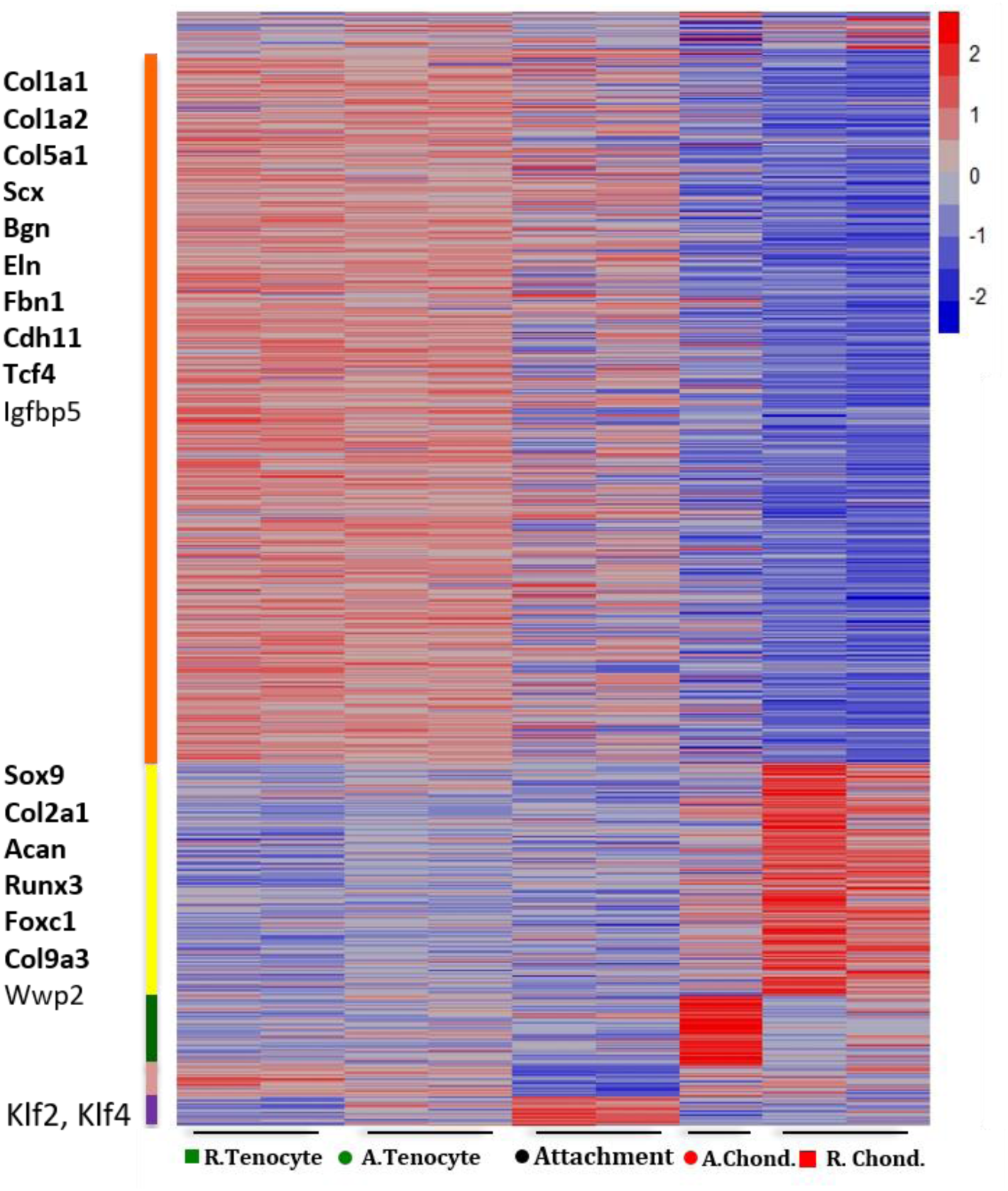
Transcriptomic analysis of tendon-to-bone attachment site domains at E14.5. Heat map of attachment site compartments, arranged along the horizontal axis of the map according to their original anatomical positions. Using CLICK, 865 genes were grouped into five clusters by expression values, which are shown after standardization. Blue-red color bar (−2-0-2) represents the log-normalized counts standardized per gene, as red is higher than the mean (0) and blue is lower than the mean. The upper cluster contains genes highly expressed in tenocytes (e.g. *Co1a1, Col1a2, Col5a1, Scx*), whereas cluster 2 contains genes highly expressed in chondrocytes (e.g. *Sox9, Col2a1, ACAN*). Cluster 3 contains genes with high expression in chondrocytes adjacent to attachment cells. Clusters 4 and 5 show genes that are low or high in attachment cells, respectively. Genes in bold are expressed in attachment cells in addition to tenocytes or chondrocytes; other genes were identified during this study.

**Figure S3:**
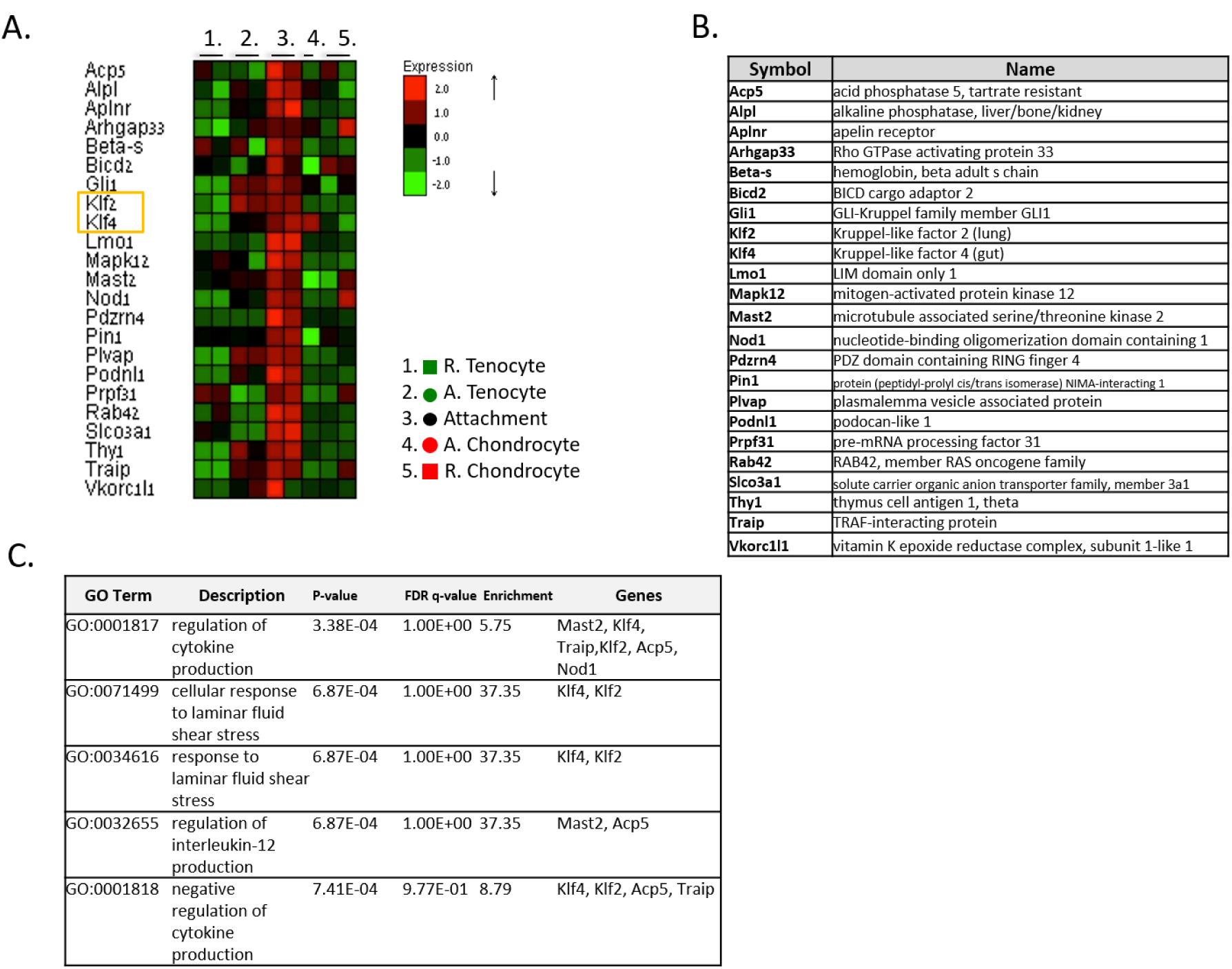
Tendon-to-bone attachment site cells up-regulated gene expression (5^th^ cluster). **(A)** Following transcriptomic analysis of attachment site domains at E14.5, the attachment site samples were ordered in the heat map in the same relative position as in the original tissue. Using CLICK, we clustered gene expression values of 865 genes into five clusters (see Materials and Methods), shown in Fig. S2. The fifth cluster contained 23 genes that were up-regulated in attachment cells. These genes, such as the KLFs, may act as regulators of the forming tendon-to-bone attachment site, and were prioritized to be extensively explored in regard to their role during tendon-to-bone attachment site development. **(B)** 5^th^ cluster list of genes, including differentiation markers such as *Thy1*, regulators of bone e.g. *Acp5* and *Alpl*, protein kinases such as *Mapk12* and *Mast2*, and signaling molecules such as *Nod, Traip, Aplnr* and others. **(C)** GO analysis of 5^th^ cluster genes.

**Figure S4:**
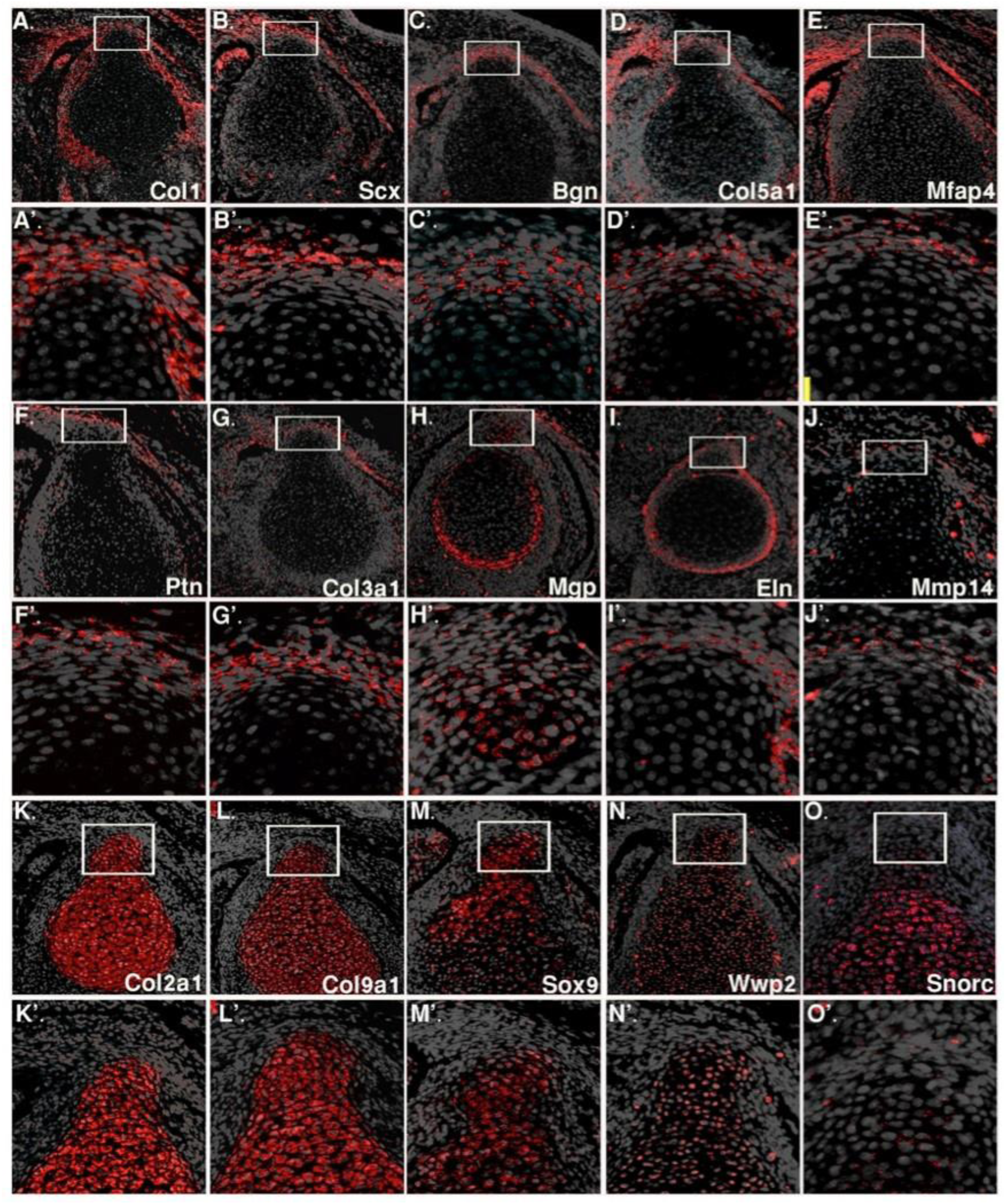
Fluorescent *in situ* hybridization analysis validated the expression profile of attachment cells revealed by RNA-Sequencing (MARS-Seq). Fluorescent *in situ* hybridization for tendon (*Col1a1, Scx, biglycan (bgn), Col5a1, Mfap4, Ptn, Col3a1, Mmp14*) and cartilage (*Col2a1, Col9a1, Sox9, Wwp2* or *Snorc*) genes at E14.5 transverse sections of the humerus, on the background of DAPI staining (gray). *In situ* hybridization for ECM genes *Mgp* and *Eln* (H,I) shows high expression in the attachment site and perichondrium, as indicated by MARS-Seq results. A-E, F-J,K-O: X20 magnification; A’-E’, F’-J’, K’-O’: magnification of upper panels.

**Figure S5:**
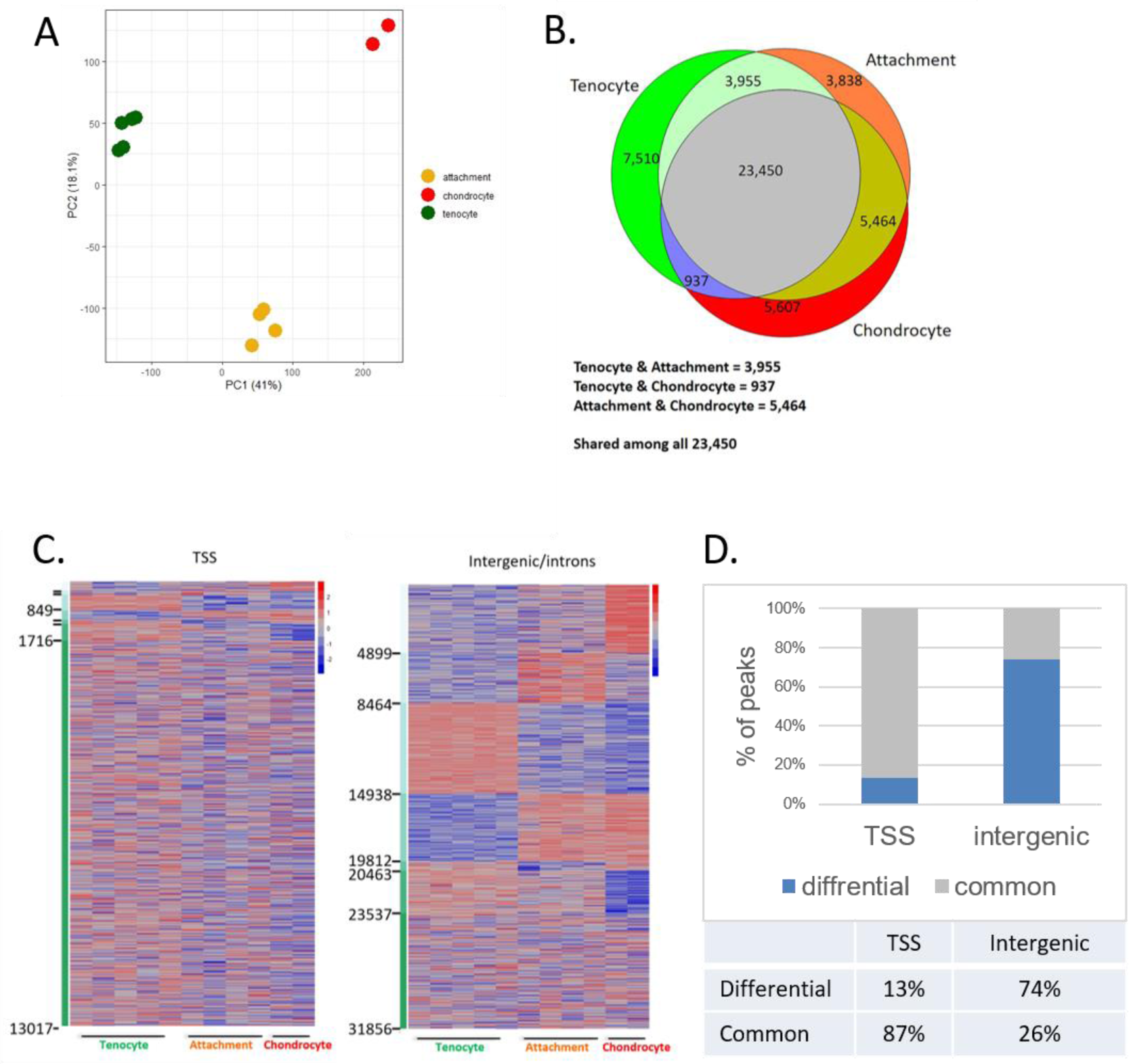
Accessible chromatin reveals an epigenetic mechanism shared by attachment cells and neighboring tenocytes or chondrocytes. **A.** PCA analysis of accessible chromatin profiles of FACS-sorted tenocytes (green), chondrocytes (red) and attachment cells (yellow). **B.** Venn diagram showing cell-specific or overlapping peaks of ATAC-Seq among tenocytes, chondrocytes and attachment cells. **C.** Heatmap of ATAC-Seq peaks. Left: TSS peaks, right: intergenic or intron peaks. **D.** Percentage of common peaks (shared by three cell types) vs. differential peaks (the chromatin is open only in one or two cell types) compared between TSS and intergenic areas.

